# Vinculin recruitment to α-catenin halts the differentiation and maturation of enterocyte progenitors to maintain homeostasis of the *Drosophila* intestine

**DOI:** 10.1101/2021.08.05.455299

**Authors:** Jérôme Bohère, Buffy L. Eldridge-Thomas, Golnar Kolahgar

## Abstract

Mechanisms communicating changes in tissue stiffness and size are particularly relevant in the intestine, because it is subject to constant mechanical stresses caused by peristalsis of its variable content. Using the *Drosophila* intestinal epithelium, we investigate the role of vinculin, one of the best characterised mechanoeffectors, which functions in both cadherin and integrin adhesion complexes. We discovered that vinculin regulates cell fate decisions, by preventing precocious activation and differentiation of intestinal progenitors into absorptive cells. It achieves this in concert with α-catenin at sites of cadherin adhesion, rather than as part of integrin function. Following asymmetric division of the stem cell into a stem cell and an enteroblast, the two cells initially remain connected by adherens junctions, where vinculin is required, only on the enteroblast side, to maintain the enteroblast in a quiescent state and inhibit further divisions of the stem cell. Removing vinculin increases enteroblast differentiation and numbers, resulting in an enlarged gut with improved ability to recover after starvation. Thus, mechanical regulation at the contact between stem cells and their progeny is used to control tissue cell number.

## Introduction

Adult epithelial tissues are maintained through the timely production of correct specialised cells by resident stem cells, ensuring tissue function and preventing disease. In the intestine for example, where the rate of tissue turnover is high, uncompensated cell loss can result in chronic inflammation, whereas over-proliferation and mis-differentiation can produce adenomas that constitute a sensitised background for tumour initiation (Gehart & Clevers, 2019). Hence, elucidating the molecular mechanisms regulating cell renewal and fate acquisition will help further our understanding of fundamental rules governing tissue homeostasis and might open avenues for therapeutic intervention.

In the mammalian intestine, a combination of locally secreted and membrane-bound proteins regulates cell fate decisions across the crypt-villus axis through the activation of key signal transduction pathways, including Wnt and Notch (Meran et al., 2017; Gehart & Clevers, 2019). There is also an increased appreciation of a potential role of mechanical cues from surrounding cells -through cell-cell and cell-matrix adhesion protein complexes-in regulating cell fate determination, as observed in other systems (Meran et al., 2017; Wickström & Niessen, 2018). Although mechanosensing of the extracellular matrix is currently under investigation in intestinal organoid cultures (Gjorevski et al., 2016), the regulation of intestinal cell fate by adhesion complexes is in general poorly understood compared to canonical signalling pathways. At adherens junctions between neighbouring epithelial cells, transmembrane E-cadherin proteins engage in homophilic adhesion via their extracellular domain, and indirectly associate with the cytoskeleton by recruitment of β-catenin via their intracellular domain, which can interact with various actin binding proteins including α-catenin. Heterodimeric transmembrane proteins integrins and their associated proteins, including talin, contribute to focal adhesions at cell-matrix junctions, thus linking extracellular ligands to cytoskeletal proteins (Sun et al., 2016).

In the intestine, as well as other mature renewing epithelia, an outstanding question is how mechanical forces are locally integrated and interpreted to fine-tune proliferation and differentiation. In *Drosophila* for example, genetic ablation of integrins in intestinal stem cells prevents overproliferation following over-activation of growth factor signalling pathways (Lin et al., 2013). However, why integrins are required for proliferation remains unsolved. Characterising *in vivo* the function of junction-associated intracellular proteins capable of responding to mechanical forces through change of conformation and activity might illuminate how mechanotransduction contributes to the regulation of tissue homeostasis.

One candidate of interest is the highly conserved actin-binding protein vinculin, as it is capable of stabilising both adherens junctions and focal adhesions under tension (Bays & DeMali, 2017) and of regulating cell-fate decisions in cell culture models and *in vivo* in the mouse skin (Holle et al., 2013; Kuroda et al., 2017; P. Wang et al., 2019; Biswas et al., 2021). Vinculin protein consists of a head and a tail domain separated by a linker region. In cells, vinculin transits from a closed, auto-inhibited inactive conformation (whereby the head and tail domains interact with each other, preventing further protein-protein interactions), to an open, active conformation initiated by force-dependent protein unfolding (reviewed in Carisey & Ballestrem, 2011; Bays & DeMali, 2017), that enables the vinculin tail domain to bind F-actin. The head domain can bind <u>v</u>inculin <u>b</u>inding <u>s</u>ites (VBS) present in other proteins (including α-catenin and talin). Notably, intracellular tension induces conformational changes in α-catenin and talin, exposing their cryptic VBS and initiating vinculin binding (Del Rio et al., 2009; Yonemura et al., 2010; Yao et al., 2014). This allows a transition of vinculin into an open conformation able to bind to the actin cytoskeleton, in turn enabling adhesion complexes to withstand higher forces in cells under tension (Le Duc et al., 2010; Dumbauld et al., 2013; Thomas et al., 2013). Thus, vinculin is one of the best characterised mechanoeffectors, as mechanical force on both talin and α-catenin is converted into differential vinculin recruitment.

*In vivo*, loss of *vinculin* is embryonic lethal in mice due to defects in neural tube closure and heart development (Xu et al., 1998). In *Drosophila*, loss of *vinculin* is homozygous viable and fertile (Alatortsev et al., 1997; Klapholz et al., 2015), precluding an essential role during embryonic development. In post-embryonic muscles, *Drosophila* Vinculin regulates actin organisation at integrin junctions (Bharadwaj et al., 2013; Green et al., 2018), and is lethal if expressed in a constitutively active form (Maartens et al., 2016).

In epithelia, *Drosophila* vinculin contributes to myosin II mediated tension sensing at cell-cell junctions through its interaction with α-catenin (Case et al., 2015; Jurado et al., 2016; Kale et al., 2018; Alégot et al., 2019;), with functions at integrin adhesion sites in epithelia remaining to be identified. Conditional *vinculin* knock-out in the mouse skin epidermis revealed a dispensable role in reinforcing newly formed adherens junctions in the stratified epidermis (Rübsam et al., 2017), and a more prominent role in maintaining stem cells in a quiescent state via stabilisation of adherens junctions and contact inhibition in the hair follicle bulge (Biswas et al., 2021). The variety of phenotypes observed so far among different tissues and organisms suggest that vinculin functions in a highly context-specific manner, and thus, characterising its contribution to tissue homeostasis in multiple systems is required to improve our understanding of mechanotransduction.

The role of vinculin in epithelial maintenance and stem cell lineage decisions is unknown in the intestine. Here, we investigate *in vivo* the role of vinculin in the maintenance of the *Drosophila* intestine, a relatively simple model of homeostatic adult epithelium used for its genetic tractability (reviewed in Miguel-Aliaga et al., 2018). In this system, resident stem cells are scattered among differentiated absorptive and secretory cells and respond to conserved signalling transduction pathways to respectively divide and differentiate. Upon cell division, intestinal stem cells (ISCs) self-renew and produce precursor cells able to differentiate into either secretory enteroendocrine cells (EEs) or large absorptive enterocytes (ECs) (Micchelli & Perrimon, 2006; Ohlstein & Spradling, 2006). *Drosophila* ISCs and their undifferentiated daughter cells, enteroblasts (EBs), are collectively called progenitor cells and express the transcription factor *escargot* (*esg*) (fig.1A). As EBs differentiate into ECs, through changes in gene expression, cells develop membrane protrusions and adopt a migratory phenotype, allowing them to move between cells; EBs also progressively increase in cell volume, resulting in an extended contact area on the basal side (Antonello et al., 2015; Rojas Villa et al., 2019). In the *Drosophila* intestine, loss of integrin-mediated adhesion has deleterious effects on stem cell maintenance and proliferation (Goulas et al., 2012; Lin et al., 2013a; Okumura et al., 2014; You et al., 2014) and E-cadherin-mediated adhesion regulates stem cell maintenance and differentiation (Choi et al., 2011; Maeda et al., 2008; Zhai et al., 2017).

We show that vinculin regulates differentiation into the absorptive cell lineage, independently of integrins at cell-matrix sites and through its association with α-catenin at cell-cell junctions.

## Results

### Vinculin has opposite effects on progenitor numbers and cell density compared to integrin and talin

In the *Drosophila* gut, complete loss of function of the ubiquitous β-PS integrin subunit (*myospheroid, mys*) progressively induces ISC loss (Lin et al., 2013a). With the goal of uncovering intracellular factors mediating integrin function in stem cells, we investigated the role of two integrin-associated mechanosensing proteins, talin and vinculin (Atherton et al., 2016; Klapholz & Brown, 2017). Talin is important for the establishment of EC polarity (Chen et al., 2018) and for stem cell maintenance (Lin et al, 2013). The role of vinculin, which is expressed ubiquitously in the gut (suppl fig.1), has not been characterised in the adult intestine. We used RNA interference to downregulate β-PS integrin, talin and vinculin in *esg-*expressing progenitor cells visualised with Green Fluorescent Protein (GFP). Downregulation of β-PS integrin and talin led to gut shortening in the posterior region (fig.1B, C) due to loss of progenitors (see reduction in GFP^+^ cells in fig.1D, E) adopting a circular shape and detaching from the basement membrane, in contrast with control cells that displayed a triangular shape (insets, fig.1D). Expression of two different validated RNAi lines against *vinculin* (*vinc*) did not impact gut length and instead led to an increase in GFP^+^ cells number and total cell density (fig.1B-F). No rounding or detachment of the progenitor cells was observed, instead cells appeared larger suggestive of accelerated differentiation (insets fig1D). In addition, *vinc* knockdown led to an increase in GFP^+^ cells and cell density (fig.1B-F). These opposite phenotypes suggested that vinculin regulates progenitor production and tissue size most likely independently of integrin/talin adhesion and prompted us to clarify the role of vinculin in intestinal tissue maintenance.

**Figure 1:**
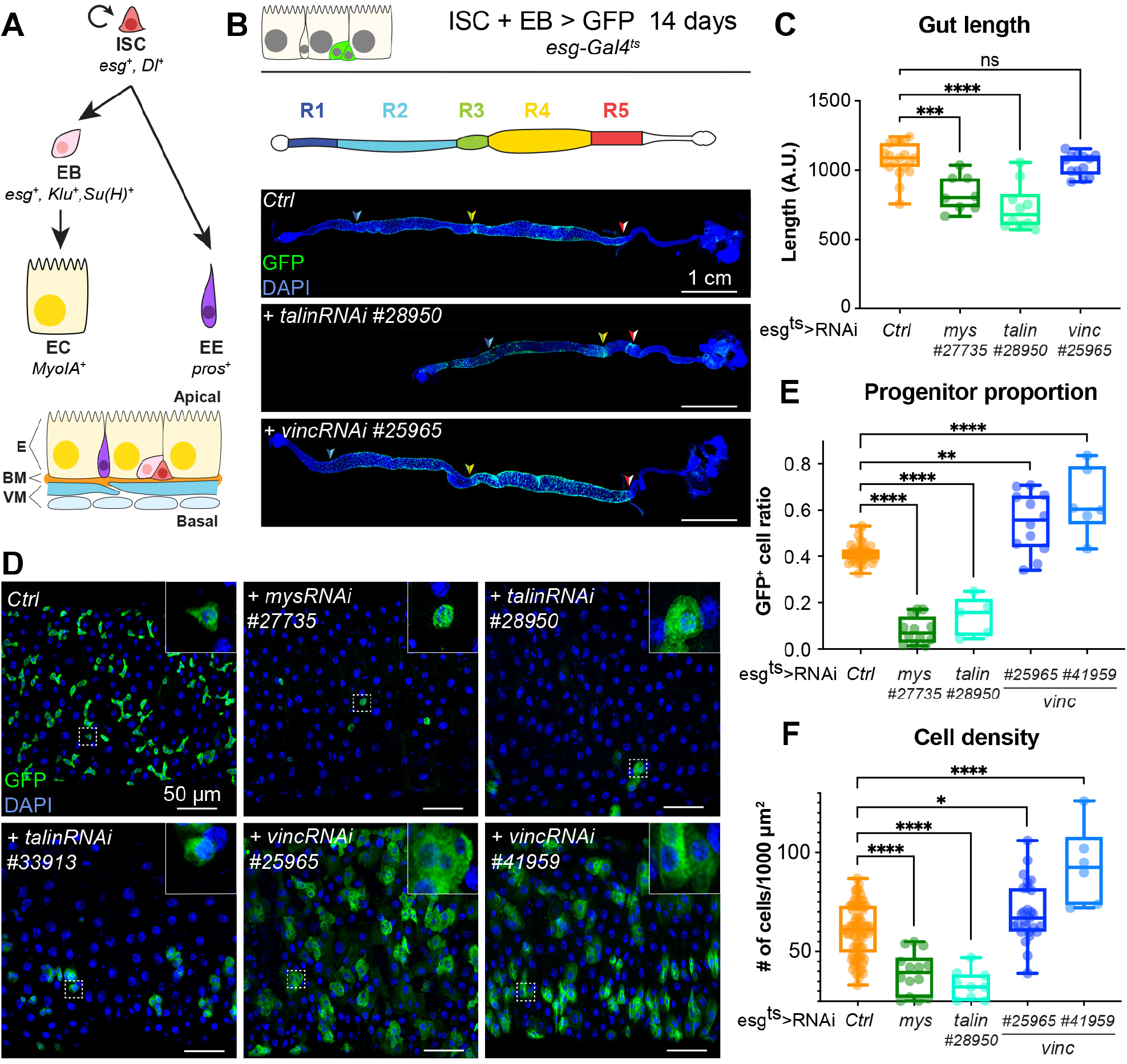
*talin* and *vinculin* knockdowns produce opposite phenotypes in the gut. **(A)** Lineage of the adult midgut: Intestinal stem cells (ISCs) self-renew and give rise to post-mitotic enteroblasts (EBs) which terminally differentiate into enterocytes (ECs), and enteroendocrine cells (EEs). Cell type-specific genes are shown in italics. Schematic at bottom shows overall tissue organisation E: Epithelium, BM: basement membrane, VM: visceral muscles. **(B, C)** Gut regions R4 and R5 are reduced by RNAi knockdown of *talin* but not *vinculin*, GFP (green) marks cells expressing the RNAi, and nuclei are blue (DAPI). Anterior is to the left in this and all subsequent figures. **(D, E)** Surface view of region R4/5. RNAi knockdown of *integrin* and *talin* shows reduced number of RNAi-expressing ISC/EB cells (GFP+) and their rounded morphology (insets). In contrast there are more ISC/EB cells in the absence of vinculin and they are enlarged. These changes result in an overall change in total cell density **(F)**. Two-tailed Mann-Whitney tests were used: ns: not significant; ****: p<0.0001, ***: p<0.001, **: p<0.01, *: p<0.05, in this and all subsequent figures.

### Intestinal cell production is increased upon global loss of *vinculin* function

To study the impact of *vinc* complete loss of function on guts, we used the null allele *vinc*^*102*.*1*^, a deletion removing approximately 30kb of genomic DNA including the entire *vinc* coding sequence (Klapholz et al., 2015). After dissection, we noted that the intestines of *vinc*^*102*.*1*^ mutants were wider compared to control *yw* flies, despite being of comparable lengths. We tested if the difference in gut width reflected a gross difference in body size by measuring the adult wing surface areas of *yw* and *vinc*^*102*.*1*^ flies. As we found them to be comparable (suppl Fig.2, and in accordance with (Sarpal et al., 2019), we next set out to establish if the increased gut width resulted from an increase in cell numbers. Guts were processed for electron microscopy (fig.2A) or immunofluorescence (fig.2B) and cells were counted in cross sections. Both methods showed increased cell numbers in *vinc*^*102*.*1*^ posterior midguts (see quantification of DAPI^+^ nuclei from cross sections of fluorescently labelled guts in fig.2B). Importantly, even when we compared guts of similar width (between 190 and 220μm in fig.2B), we found that *vinc*^*102*.*1*^ gut cross-sections had approximately twice as many cells as *yw* guts.

To characterise how vinculin regulates tissue homeostasis, we genetically labelled all intestinal progenitor cells (*esg*^+^) and their progeny with GFP (fig.2C-E), combining the labelling cassette *esg*^*ts*^*F/O* (or “escargot-Flip-Out”, Jiang et al., 2009) in *yw* or *vinc*^*102*.*1*^ male flies (see suppl fig.3A and Methods for experimental details). We compared the extent of GFP labelling as a proxy for cell production over a period of 14 days post induction. During this period, moderate tissue turn-over occurs in control *yw* guts (Jiang et al, 2009), as evidenced by groups of more than two cells that included some ECs, recognised by their large polyploid nuclei (fig.2D, E, *yw* guts). Whereas the distribution of newly produced cells was largely uniform and low across the posterior midguts of *yw* flies (fig.2D, top panel), guts of *vinc*^*102*.*1*^ flies presented with a strong labelling at the interface between R4/5 regions of the posterior midgut (regions defined in Buchon et al., 2013) (fig.2D middle panel, compare to cartoon fig 1B). Quantification of GFP signal intensity (as described in suppl fig.2B) across 27 guts showed a penetrant phenotype (see heat maps fig.2D). The phenotype was fully rescued in flies carrying a genomic *Vinc* rescue construct (Vinc-RFP, Klapholz et al., 2015) (n=34, fig.2D, bottom panel), confirming that it is *vinc* loss that accelerates tissue turnover. This was confirmed by measuring gut width (fig.2F) and cell density (fig.2G), which both reflected increased cell production. Altogether, the increase in cells observed when vinculin is removed indicates that vinculin inhibits cell renewal under basal conditions.

**Figure 2:**
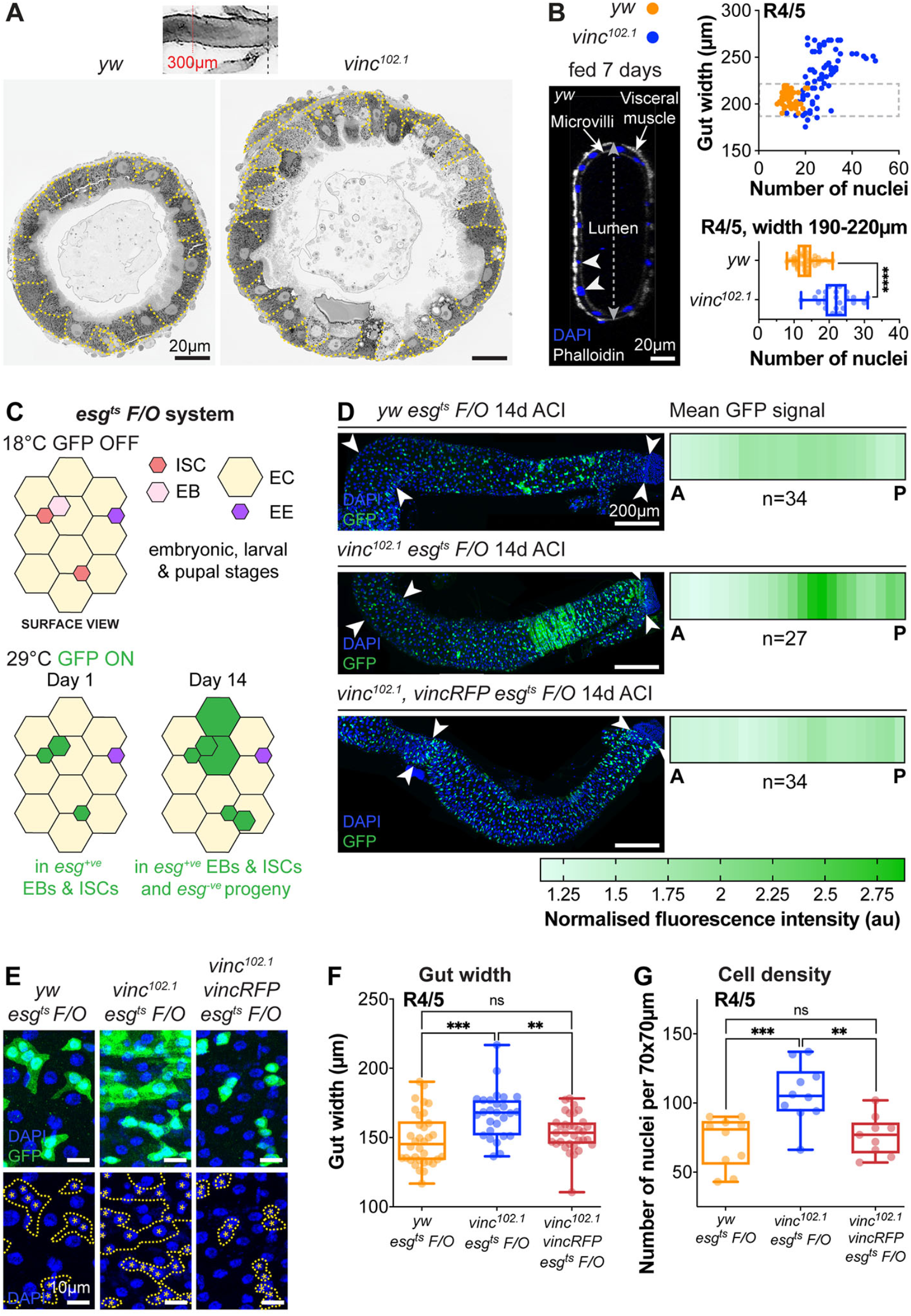
Global loss of vinculin accelerates intestinal cell production. **(A)** SEM cross-sections of *yw* and *vinc*^*102*.*1*^ midguts. Dashed lines indicate cell boundaries. *vinc*^*102*.*1*^ guts are wider than *yw* guts. (**B)** Confocal cross-sections of midguts stained with Phalloidin (white) and DAPI (blue) were analysed to extract gut width (arrow in lumen) and epithelial cell numbers (arrowheads indicate nuclei of epithelial cells). Nuclei number per gut cross section were plotted against R4/5 midgut width. *yw*: n = 55 datapoints from five guts, *vinc*^*102*.*1*^: n= 77 datapoints from seven guts. *vinc*^*102*.*1*^ midguts contain more cells than *yw*, even when comparing guts of the same width. (subset of guts with comparable width (dashed box) replotted in bottom graph *yw*: n= 55 datapoints from five guts, *vinc*^*102*.*1*^: n= 34 datapoints from four guts.) **(C)** Top view schematic of the *esg*^*ts*^ *F/O* system. At the permissive temperature (29°C), the ISC- and EB-specific *esg-Gal4* drives expression of both *UAS-GFP* and *UAS-flp*, which mediates permanent, heritable expression of GFP. Thus, as ISCs divide and EBs differentiate during adulthood, all cells which arise from progenitors will express GFP. **(D)** Left panels: Progenitors and newly produced cells (GFP^+^, green) 14 days after clone induction (ACI). A large GFP^+^ region is present in the R4/5 region of *vinc*^*102*.*1*^ *esg*^*ts*^ *F/O* guts. Arrowheads indicate the region along which heatmaps were generated. Images are z projections through half the gut depth. Right panels: heatmaps of GFP-fluorescence intensity. Dark green corresponds to high levels of GFP fluorescence, indicating elevated tissue turnover. *yw esg*^*ts*^ *F/O* n=34 guts, *vinc*^*102*.*1*^ *esg*^*ts*^ *F/O* n=27 guts, *vinc*^*102*.*1*^ *vincRFP esg*^*ts*^ *F/O* n=34 guts, from three replicates. **(E)** *esg*^*ts*^ *F/O* clones with GFP-DAPI overlay and DAPI only. Dashed lines indicate clone boundaries. Asterisks indicate GFP^+^ nuclei. Images are z-projections. **(F)** Quantification of gut width. *yw esg*^*ts*^ *F/O* n= 34 guts, *vinc*^*102*.*1*^ *esg*^*ts*^ *F/O* n= 27 guts, *vinc*^*102*.*1*^ *vincRFP esg*^*ts*^ *F/O* n=34 guts. **(G)** Quantification of cell density. *yw esg*^*ts*^ *F/O* n= 10 guts, *vinc*^*102*.*1*^ *esg*^*ts*^ *F/O* n= 10 guts, *vinc*^*102*.*1*^ *vincRFP esg*^*ts*^ *F/O* n=9 guts

### Vinculin is not required in intestinal stem cells to regulate proliferation

As *Drosophila* intestinal cell production relies solely on ISCs (Micchelli & Perrimon, 2006; Ohlstein & Spradling, 2006), we speculated that vinculin might negatively regulate stem cell proliferation, in a similar fashion as the recently observed control of bulge stem cell proliferation in mice hair follicle (Biswas et al., 2021).

**Figure 3:**
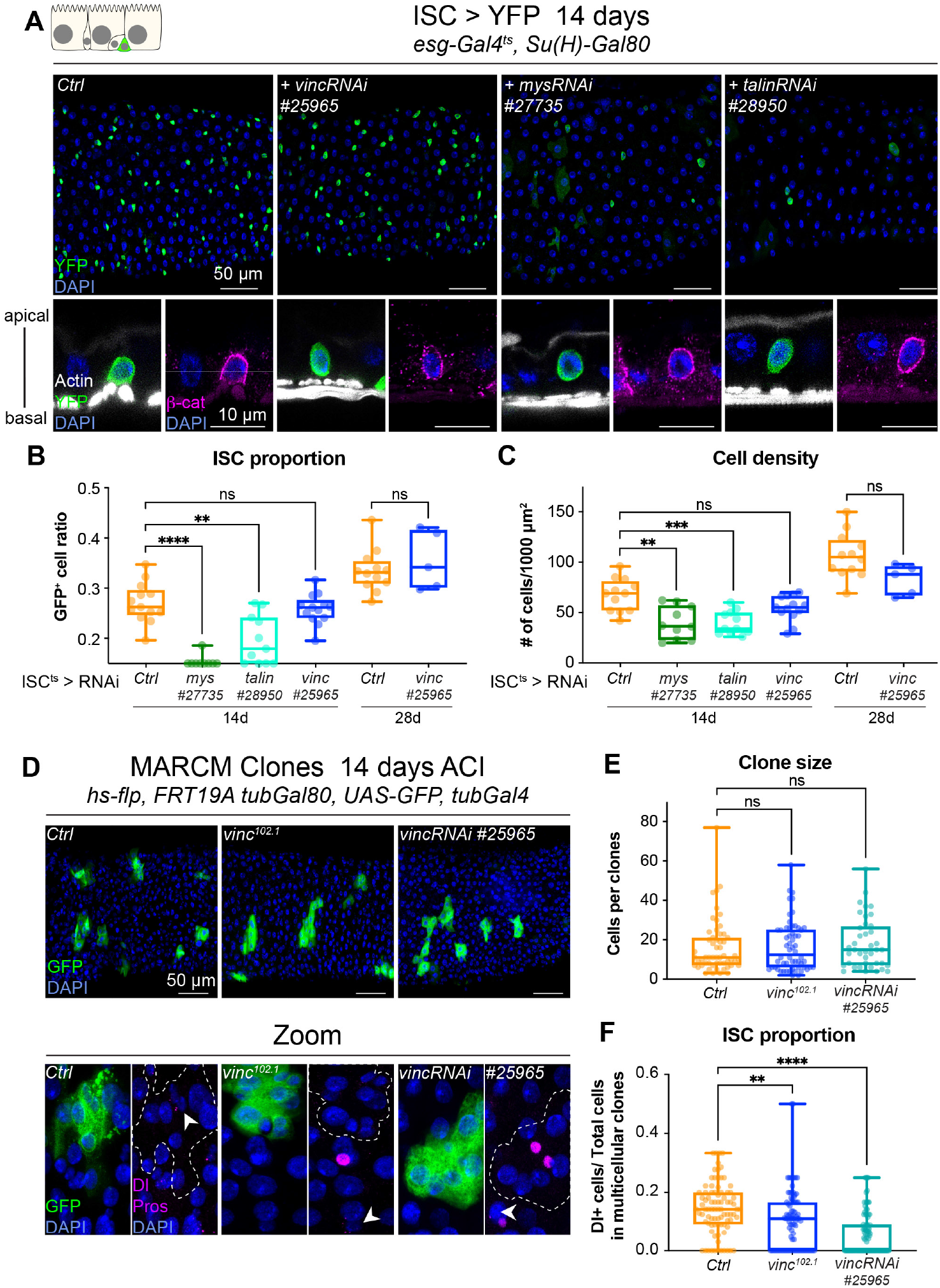
Vinculin is not required in stem cells to regulate cell proliferation. **(A)** 14-day RNAi-mediated knockdown of *mys, talin* and *vinculin* in ISCs (GFP, bright green). Nuclei are blue throughout the figure (DAPI). *mys* and *talin*, but not *vinc*, knockdown induce stem cell rounding and detachment from the basal side (orthogonal views from guts stained with Phalloidin (white) and β-catenin (β-cat, magenta) **(B-C)** Quantification of the ratio of GFP^+^ ISCs to the total number of cells (B) or cell density (C), after 14 or 28 days of RNAi expression. **(D)** GFP-labelled mitotic clones representing wild-type (left), *vinc*^*102*.*1*^ homozygous mutant (middle) or *vinc RNAi* expressing cells (right). In all conditions, unlabelled cells are considered wild-type. Zoom panels show clones (GFP, green) outlined with dotted lines, ISCs (anti-Dl, arrowheads, magenta), EEs (anti-Pros, magenta nuclear signal). Examples of multicellular clones devoid of Dl staining are shown for *vinc*^*102*.*1*^ and *vinc RNAi*. **(E)** Quantification of cell numbers in clones containing Dl^+^ cells. Ctrl: n=56 clones across 26 guts; *vinc*^*102*.*1*^: n=58 clones across 34 guts; *VincRNAi:* n=40 clones across 25 guts. **(F)** Quantification of stem cell proportion in multicellular clones (number of Dl^+^ cells/number of cells per clone). Ctrl: n=85 clones across 26 guts; *vinc*^*102*.*1*^: n=86 clones across 34 guts; *VincRNAi:* n=92 clones across 25 guts.

We expressed *vinc* RNAi and YFP in all ISCs and monitored the stem cell pool and tissue size after 14 and 28 days of expression at permissive temperature (fig.3A-C). Unlike *vinc* knockdown in both ISCs and EBs, we did not detect any change in stem cell proportion or tissue density (fig.3B-C). This was again in contrast to *mys* or *talin* knockdown that led to a clear rounding and detachment of ISCs from the basement membrane as seen by the delocalisation of β-Catenin to the basal side of the cell (fig.3A,orthogonal views), and possibly explaining the significant reduction of the stem cell population (fig.3B) and cell density (fig.3C) not observed with ISC-specific *vinc* knockdown.

As an alternative way to test the role of vinculin in stem cells, we generated GFP-labelled control, homozygous *vinc*^*102*.*1*^ mutant or *vinc*-RNAi expressing mitotic clones derived from a single stem cell and compared the size and composition of multicellular clones in tissues stained for Delta (Dl, ISCs) and Prospero (Pros, EEs). Clones of all genotypes containing at least one Dl^+^ ISC had comparable cell numbers 14 days after clone induction (fig.3D-E), suggesting comparable rates of cell production, and thus no effect on basal rates of proliferation. Additionally, we did not notice any change in EE proportions (suppl fig.4A), indicating that loss of *vinc* in ISCs does not affect differentiation of secretory cells. We noticed however that the proportion of Dl^+^ ISCs was reduced in *vinc*^*102*.*1*^ mutant or *vinc*-RNAi expressing clones (fig.3F), with some clones devoid of Dl staining (see Zooms in fig.3E) but containing numerous EBs (recognised as Pros^-^ small cells, suppl fig.4B), suggesting that vinculin might control EB differentiation. Altogether, these data led us to conclude that vinculin does not have a role in regulating stem cell proliferation, so that is not the cause of the increased numbers of cells when vinculin is specifically absent in this cell type.

### Vinculin maintains enteroblasts in a quiescent state

To examine the role of vinculin in EBs, we first sought to examine the EB pool when *vinc* was depleted. Notch is activated in EBs (fig.1A) (Micchelli & Perrimon, 2006; Ohlstein & Spradling, 2006, 2007), thus we combined *yw* or *vinc*^*102*.*1*^ males to a <u>N</u>otch <u>R</u>esponsive <u>E</u>lement fused to lacZ (NRE-lacZ, Furriols & Bray, 2001) and immunostained dissected guts for β-Galactosidase and Dl to quantify the relative proportions of EBs and ISCs respectively (fig.4A-C). *vinc*^*102*.*1*^ guts had a clear accumulation of NRE-lacZ^+^ cells (especially in the region of high turnover R4/5 (marked with longitudinal arrows in the *vinc*^*102*.*1*^ gut in fig.4A), but the overall proportion of stem cells was reduced (fig.4C). We had a similar observation using a transcriptional reporter of the JAK-STAT signalling pathway, which is also required during enterocyte differentiation (Beebe et al., 2010) (suppl fig.5).

It was somewhat surprising to observe cells with active Notch signalling accumulating in a context where the ligand-presenting cells (Dl^+^ ISCs) were sparse. One possible explanation would be that instead of dividing predominantly asymmetrically to produce one stem cell and one daughter EB cell with Notch signalling active, as described in homeostasis (O’Brien et al., 2011; De Navascués et al., 2012; Guisoni et al., 2017; Hu & Jasper, 2019), *vinc*^*102*.*1*^ ISCs may more frequently divide symmetrically, producing two EBs with Notch signalling active. This would be compatible with the observed reduction of ISC ratio in mitotic clones (fig.3F).

Another possibility would be that EBs progress faster down the differentiation path. Indeed, under normal conditions, diploid EBs remain dormant after production, until in response to tissue demand, they activate signalling programmes leading to their maturation, migration, endoreplication and differentiation into ECs (Antonello et al., 2015; Choi et al., 2011; Rojas Villa et al., 2019; Xiang et al., 2017; Zhai et al., 2017).

To distinguish between dormant and activated EBs, we measured the nuclear size of ISCs (Dl^+^), EBs (*NRE-lacZ*^+^), and ECs (Dl^-^*NRE-lacZ*^*-*^ and polyploid), as a proxy of the degree of differentiation, since differentiation involves the cells becoming increasingly polyploid (fig.4D). In control tissues, EBs were distributed in two groups (see *yw* violin plots, fig.4D), with dormant and activated cells corresponding to smaller and bigger nuclei/cells respectively (Antonello et al., 2015; Rojas Villa et al., 2019). In contrast, the large majority of *vinc*^*102*.*1*^ EBs had bigger nuclei, suggesting reduced numbers of cells were in the dormant stage in the absence of *vinc*. We also noted that most ECs in *vinc*^*102*.*1*^ guts had smaller nuclei compared to *yw* guts, suggesting incomplete maturation of ECs (fig.4D).

**Figure 4:**
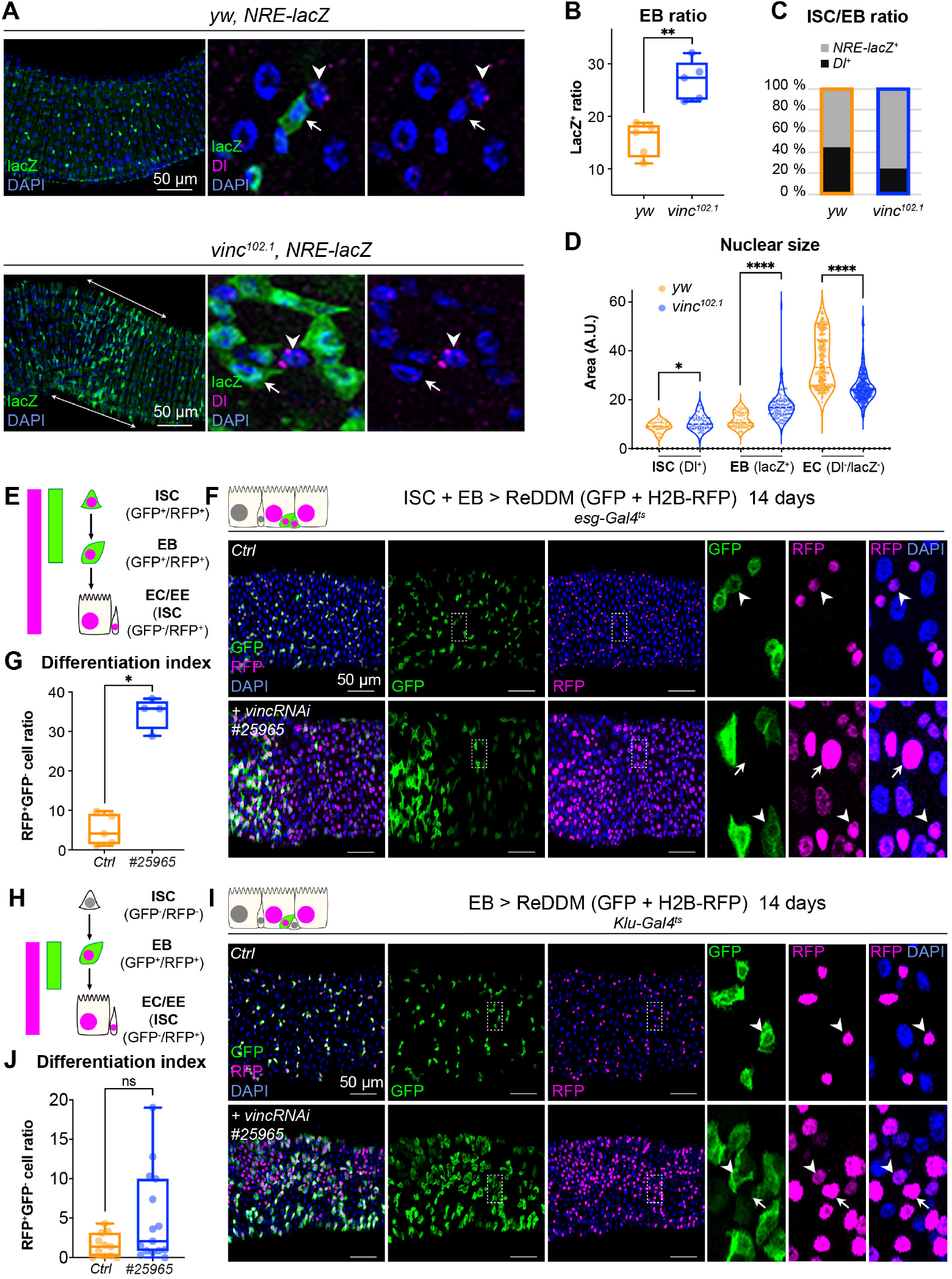
Vinculin slows enterocyte precursor differentiation and tissue turnover. **(A)** EBs in 14-day-old *yw* or *vinc*^*102*.*1*^ guts of male flies, marked by expression of Notch reporter NRE-lacZ (β-galactosidase antibody staining green, arrows in insets). Dl marks ISCs (magenta, arrowheads). Nuclei are blue throughout the figure (DAPI). Arrows highlight the region R4/5 where NRE-lacZ^+^ cells accumulate in *vinc*^*102*.*1*^ guts. **(B)** Quantification of EB ratio (number of NRE-lacZ^+^ cells/ total number of cells) showing the excess of EBs in *vinc*^*102*.*1*^ guts. **(C)** Quantification of ISC/EB ratio. **(D)** Quantification of nuclear size of the different cell populations. Note the distribution of EBs in two groups in *yw* guts, representing dormant and activated EBs. *vinc*^*102*.*1*^ EBs have overall larger nuclear size. **(E)** Schematic of the ReDDM system expressed in progenitor cells. **(F)** Expression of ReDDM in progenitors with *vincRNAi*^*#25965*^ or without (Ctrl) for 14 days. Close ups (dashed line rectangles) show GFP^-^RFP^+^ differentiated cells (arrows) upon *vinc* knockdown. Arrowheads indicate GFP^+^RFP^+^ progenitors. **(G)** Quantification of the differentiation index (proportion of GFP^-^ RFP^+^ cells) Expression of *vincRNAi*^*#25965*^ in progenitors accelerates differentiation. **(H)** Schematic of the ReDDM system expressed in EBs. **(I)** *vincRNAi*^*#25965*^ expression in EBs leads to the accumulation of GFP^+^ cells. Close ups show GFP^+^RFP^+^ (arrowheads) and GFP^-^RFP^+^ (arrows) cells, the latter being more frequent in *vincRNAi*^*#25965*^ *expressing guts*. **(J)** Quantification of the differentiation index when RedDM is expressed with *Klu-Gal4*. No statistical difference was noted when *vincRNAi*^*#25965*^ was expressed, suggesting Klu^+^ cells accumulate and stall in this condition.

To evaluate the role of vinculin in EBs, we compared loss of vinculin in both ISCs and EBs, (*esg*^*+*^*>VincRNAi*) versus EBs only (*klu*^*+*^*>VincRNAi*, Korzelius et al., 2019; Reiff et al., 2019). To evaluate how vinculin controls the rate of EB differentiation, we expressed under the control of the same promoter two fluorophores, GFP and RFP, with differing protein half-lives (ReDDM, Antonello et al., 2015), such that newly differentiated EBs can be recognised by the absence of GFP and presence of nuclear RFP (see methods and cartoons in fig.4E, H). *vinc* RNAi expression in *esg*^+^ cells led to faster progenitor differentiation cells (fig.4E, F; note the accumulation of GFP^-^RFP^+^ cells) and an increased differentiation index calculated as the ratio of GFP^-^ RFP^+^ cells/total cells (fig.4G). Expression of *vinc RNAi* in *klu*^*+*^ cells instead led to a sharp increase of the proportion of GFP^+^ EBs (fig.4H, I, suppl fig.6D), despite no significant rise in the differentiation index (fig.4J). In the later stages of EB differentiation, the Notch responsive Klu transcription factor represses the secretory fate in EBs and contributes to their switch to endoreplication as they differentiate into ECs (Korzelius et al., 2019). Our observation that cells expressing *klu* accumulated upon *vinc* RNAi expression suggested EBs started to differentiate but did not complete their maturation as Klu should be turned off in ECs (Korzelius et al., 2019). The absence of terminal EB differentiation could thus be due to misregulation of *klu* or lack of tissue demand for new absorptive cells. In addition, compensatory ISC proliferation contributes to the accumulation of Klu^+^ cells, as we observed an increase in mitotic cells (PH3^+^) in posterior midguts of *klu-Gal4*>*vinc* RNAi guts (suppl fig.6A-C). This suggests that specific loss of vinculin in EBs feeds back on ISCs, stimulating them to divide. Occasionally, we observed instances of mitotic EBs (PH3^+^ Klu^+^ cells) following *vinc* RNAi expression (suppl fig.6B). This abnormal occurrence of mitotic marker in EB cells, which are normally post-mitotic, suggests these newly generated EBs have kept some mitotic potential, fitting with an incomplete maturation due to accelerated differentiation upon loss of *vinc*. Indeed, these cells are reminiscent of the division capable EB-like cells previously described (Kohlmaier et al., 2015). Finally, *vinc* knockdown in ECs did not elicit an increased cell density (suppl fig.7), so we conclude that vinculin acts specifically in EBs to maintain them in a dormant state to tune differentiation rates to tissue needs.

### Vinculin intestinal function is mediated by its association with α-catenin

Adherens junctions between dormant EBs and ISCs regulate ISC proliferation (Choi et al., 2011; Zhai et al., 2017). Therefore, a function of vinculin that would explain its role in maintaining EB quiescence and reduced ISC proliferation might be via regulation of these junctions. We found that vinculin was present in these junctions, co-localising with α-catenin (suppl fig.1E), and consistent with being downstream of α-catenin, knockdown of *vinc* did not alter α-catenin membrane localisation (suppl fig.8A).

If the effects of loss of vinculin in EBs are via its binding to α-catenin, then we may expect similar defects when α-catenin is removed. We therefore examined the phenotype of *α-catenin (α-cat)* downregulation in *esg*^*+*^ progenitor cells (fig.5A-C). Nuclear size quantification of GFP^+^ progenitor cells showed that as in *vinc* RNAi expressing guts, *α-cat* RNAi expression resulted in larger *esg+* cells, equivalent in size to differentiating EBs and ECs (fig.5B) and an overall accumulation of EBs (fig.5C). Thus, α-catenin is required in progenitor cells to regulate tissue turnover.

To test whether vinculin function was mediated by its ability to bind to α-catenin via the VBS domain, we downregulated *α-cat* as above and concomitantly expressed RNAi-resistant α-catenin constructs containing or lacking domains impacting vinculin binding (Alégot et al., 2019) (fig.5D). As before we used the ReDDM lineage timer to measure the degree of differentiation (fig.5E). Guts rescued by co-expression of full-length α-catenin (α−Cat FL) were indistinguishable from wild-type controls (expression of ReDDM only), with similar rates of differentiation (fig.5E-G). In contrast, co-expression of α-catenin constructs with the VBS deleted (α−Cat ΔM1a and ΔM1b), failed to rescue EBs accumulation. Deleting the adjacent M2 domain did not impair the ability of α-catenin to rescue this phenotype (fig 5E-G).

**Figure 5:**
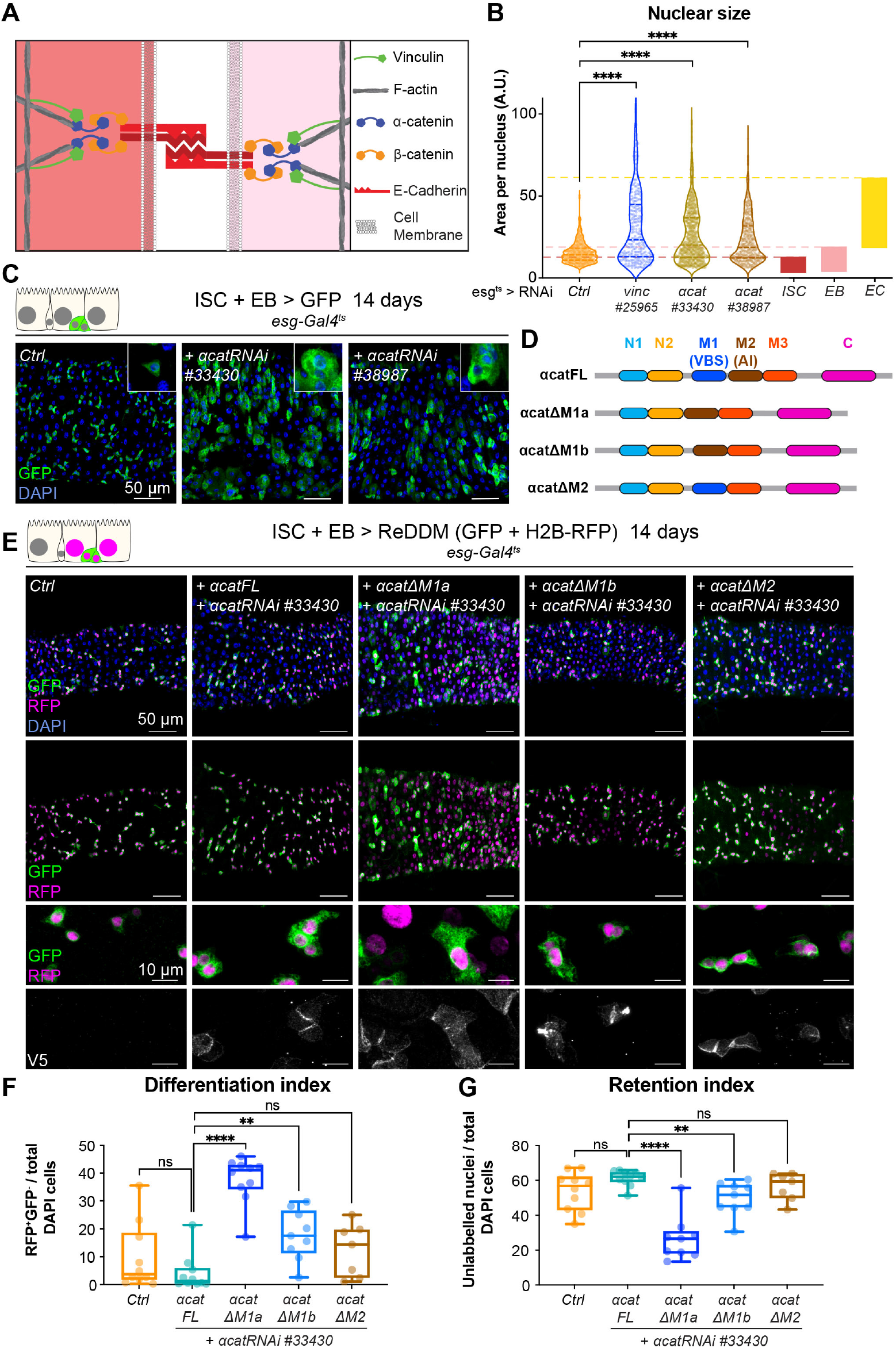
Vinculin recruitment to α-catenin prevents enterocyte differentiation. **(A)** Cartoon depicting activated vinculin binding to α-catenin and actin filaments at adherens junctions. **(B-C)** Expression of two different *α-cat* RNAi in progenitors (marked by GFP, green) for 14 days. **(B)** Quantification of nuclear size in GFP^+^ cells for indicated genotypes. Cells were recorded from a minimum of 3 guts per genotype. The coloured boxes ISC, EB, EC on the right-hand side of the graph indicate nuclear size distributions of each cell type determined in *yw* guts (as in fig. 4D). Expression of *vinc* or *α-cat* RNAi produce an accumulation of cells with nuclear size equivalent to large EBs and ECs. **(C)** Representative images of guts analysed in B. The insets show clusters of GFP cells, comparable to those observed when *vinc* RNAi is expressed (see Fig. 1D). **(D)** Summary of the rescuing α-catenin constructs used. FL: full length α-cat; ΔX: full length depleted of indicated domain. The different α-cat domains are coloured. M1 contains the vinculin binding site (VBS). M2 contains the auto-inhibitory domain. **(E)** Expression of ReDDM in progenitors with α*-cat RNAi*^*#33430*^ and various V5-tagged α-cat rescuing constructs for 14 days. Ctrl represent expression of ReDDM only. Nuclei are blue (DAPI). High magnification panels show V5 staining (white), showing accumulation of α-catenin at cell junctions in all conditions, and cytoplasmic enrichment of the ΔM1a fusion protein. **(F)** Quantification of the differentiation index (proportion of newly differentiated GFP^-^RFP^+^ cells) indicates that expression of α-cat ΔM1a in *α-cat* RNAi accelerates differentiation. **(G)** Quantification of the proportion of unlabelled cells indicates the cell retention rate. Expression of α-cat ΔM1a in *α-cat* RNAi accelerates tissue turnover.

Clearly, vinculin represses activation of a pool of EC precursors via its interaction with α-catenin. This suggests a model where vinculin, in its open, active form, contributes to keeping cellular junctions of EBs under tension, and this keeps the cells in a dormant state. In the absence of vinculin (or as the protein switches back to a closed, inactive conformation), EB junctions may destabilise faster, facilitating the transition from a dormant to an activated, migratory EB state and further differentiation into ECs.

We reasoned that if this model is correct then: (i) expressing a constitutively open form of vinculin (Vinc^CO^, Maartens et al., 2016) should rescue the accelerated turnover of *vinc*^*102*.*1*^ mutants; (ii) increasing cortical tension in progenitor cells should prevent them from adopting the EC fate; and (iii) decreasing cortical tension should accelerate tissue turnover. To test if vinculin function in homeostasis depends on the open active form, we compared *vinc*^*102*.*1*^; *esg*^*ts*^ _*F/O* guts (as in fig.2C, D) to similar guts simultaneously expressing *UAS-vinc*^*CO*^ in all GFP cells. We observed that expressing the active form of vinculin led to a partial rescue of the *vinc*^*102*.*1*^ phenotype, as we no longer observed large patches of newly generated GFP^+^ cells (suppl fig.9). To alter the contractility of progenitor cells, we modulated myosin II activity. *Drosophila* myosin II is composed of two regulatory light chains and a heavy chain, encoded respectively by *spaghetti-squash* (*sqh*), *myosin light chain-cytoplasmic* (*mlc-c*) and *zipper* (*zip*) (Franke et al., 2006). The activity of myosin II can be elevated experimentally by expression of a phospho-mimetic, active, form of Sqh (Sqh^DD^, Mitonaka et al., 2007). Expression for 14 days resulted in small round progenitor cells (similar to dormant EBs described in Rojas Villa et al., 2019), that remained undifferentiated. Instead, RNAi-mediated downregulation of *sqh* or *zip* (predicted to reduce cell contractility) mildly increased differentiation rates, and many EBs cells changed shape and were visibly more advanced towards EC differentiation (suppl fig.10), reminiscent of the semi-differentiated status of *vinc* RNAi expressing EBs.

Altogether, our data supports a model whereby vinculin associates with the VBS of α-catenin in EBs, which becomes exposed upon force-induced α-catenin conformational change. This binding prevents premature EB differentiation and compensatory ISC proliferation. The idea that vinculin functions by mediating increased tension on adherens junction was supported by the effects of independently altering acto-myosin function.

### Vinculin mutant flies are more resilient to starvation

Our experiments have focussed on the gut in a homeostatic state, however plasticity is an important component of intestinal function with stem cell proliferation and progenitor differentiation able to transiently alter in response to diet and regenerate tissue after damage, before reverting to steady-state levels (e,g. Buchon et al., 2010; O’Brien et al., 2011). Since vinculin has a role in preventing premature EB differentiation and compensatory ISC proliferation, we hypothesised that *vinc* mutants might display perturbed intestinal regeneration.

To test this, we subjected flies to a cycle of starvation and refeeding as starvation results in cell depletion and gut shortening, whilst refeeding induces cell proliferation and gut growth (McLeod et al., 2010; O’Brien et al., 2011; Lucchetta & Ohlstein, 2017). Cohorts of mated *yw* or *vinc*^*102*.*1*^ females were either continuously fed on cornmeal food for the duration of the experiment or fed for 7 days, starved for 7 days (provided with water only) and then refed for a maximum of 14 days. Guts were dissected at various time points (fig.6A,B), and gut length and width measured. We observed expected reversible changes in gut length (suppl fig.9) and width (fig.6C) for both genotypes, indicating that vinculin is dispensable for diet-induced tissue clearance and repopulation.

Two differences between *yw* and *vinc*^*102*.*1*^ flies were however noted. First, following a period of refeeding after starvation, the width of *vinc*^*102*.*1*^, but not *yw*, midguts initially exceeded the original size before eventually returning to it (compare days 14 and 18, fig 6D), suggestive of an excess in EB activation and differentiation. Second, *vinc*^*102*.*1*^ flies showed better survival to the starvation regime compared to *yw* flies (fig.6G). Possible reasons for this phenotype could be that *vinc*^*102*.*1*^ guts started off with a larger number of progenitor cells, which could have minimised the deleterious effects of cell loss and/or provided a greater pool of cells ready to start differentiating in the right nutritional conditions.

**Figure 6:**
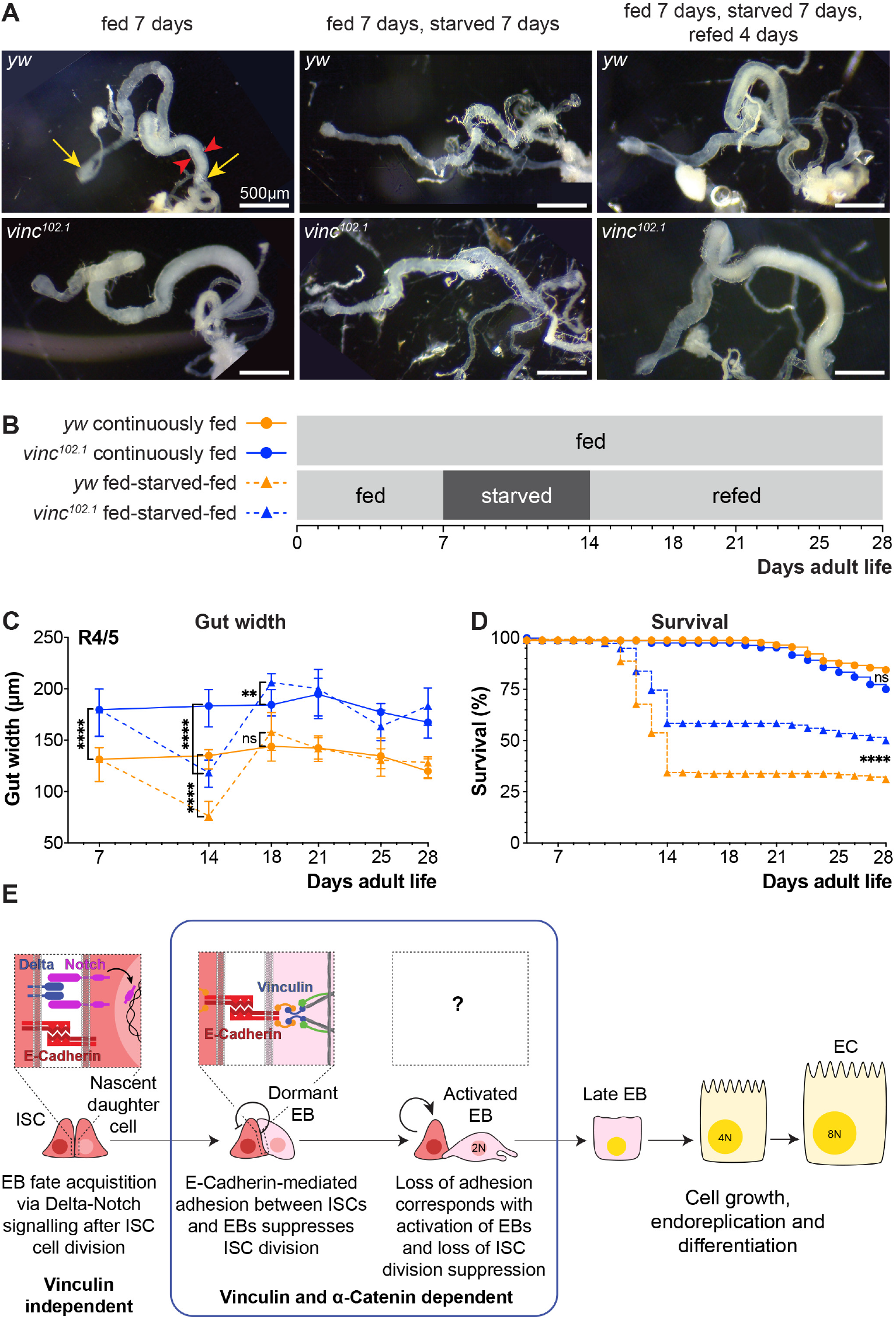
Loss of vinculin activity promotes tissue regeneration and organismal fitness. (A) Gross midgut size changes during feeding, starving and refeeding. Gut width was recorded at the R4/5 position (arrowheads). **(B)** Schematic of the feeding regime. Flies from each genotype and condition were dissected and midgut length and width measured at indicated days. **(C)** Following 7 days of starvation, R4/5 midgut width is significantly narrower in fed-starved compared to continuously fed flies for both *yw* and *vinc*^*102*.*1*^. *yw* midgut width recovers after four days of refeeding, however, *vinc*^*102*.*1*^ midgut width of fed-starved-refed flies exceeds that of continuously fed flies at this time point, before reverting back to original gut width. Symbols represent the median of 8-23 midguts measured, error bars indicate interquartile range. **(D)** Survival during cycle of feeding, starving and refeeding as shown in (B). After the third day of starvation considerable death is experienced by both *yw* and *vinc*^*102*.*1*^ flies, with *yw* showing the poorest survival. After transfer to refed conditions, remaining starved flies survive well. **(E)** Model: reinforcement of cell adhesion by α-catenin and vinculin helps to maintain EBs in a dormant state. The mechanical or chemical cues that relieve vinculin from the adhesion complex and promote tissue renewal remain to be identified.

These experiments lead us to propose a model whereby vinculin functions within EBs to maintain their dormancy and prevent adjacent ISCs from proliferating, most likely by increasing the force on cell junctions (fig.6E). The mechanical or chemical cues that relieve vinculin from the adhesion complex and promote tissue renewal remain to be identified.

## Discussion

Recent *in vitro* manipulation of tissues, cells and substrates have demonstrated the importance of mechanotransduction in cell fate specification (Kumar et al., 2017). Here, we employed genetic approaches to establish *in vivo* the role of the conserved mechanoeffector protein vinculin in intestinal tissue homeostasis.

### Vinculin dependent junctional stability regulates intestinal tissue turnover

Using the *Drosophila* intestine as a simple model of an established self-renewing adult epithelium, we have discovered that vinculin regulates EB differentiation. With conditional and cell-specific knockdowns, we demonstrated that unlike two core members of cell-matrix adhesion complex, the integrin βPS subunit and talin, which directly binds vinculin, vinculin appears dispensable for intestinal stem cell maintenance. This was surprising to us given the well-known role of vinculin in focal adhesion stabilisation *in vitro* (reviewed in Carisey & Ballestrem, 2011; Bays & DeMali, 2017). Our work instead showed that vinculin depletion in EBs induces their partial maturation towards absorptive cells, in turn accelerating tissue turnover. As this function is dependent on interaction with α-catenin, we concluded that vinculin may be required in EBs to stabilise adherens junctions. As the *vinc* phenotype correlated with that caused by decreased levels of myosin II activity, it is likely that in the intestine vinculin senses myosin II tension, as in other developing epithelia in mammals (Rübsam et al., 2017) and *Drosophila* (Case et al., 2015; Jurado et al., 2016; Kale et al., 2018; Alégot et al., 2019). This could explain why in ISC/EB pairs, vinculin is only required on the EB side where cytoskeletal tension vary as cells undergo morphological changes during maturation and differentiation. Recent work in cell culture models showed that vinculin can be recruited asymmetrically in a force-dependent manner at sites of cadherin adhesion near mitotic cells. Whilst in this case, asymmetric recruitment promotes cell shape changes during cell division and maintenance of epithelial integrity (Monster et al, 2021), our study reveals that asymmetric vinculin requirements at cell junctions can also regulate cell fate decisions and tissue composition in the intestine. Our findings place vinculin in a feedback loop that couples the process of enterocyte differentiation to stem cell proliferation in the intestine. In the absence of vinculin, intestinal stem cells constitutively sense a demand for tissue replenishment and produce new cells, despite the tissue containing high proportions of progenitors, thus generating increased cell density.

Previous work showed that in response to poor nutritional conditions (or insulin signalling pathway impairment), adherens junctions between ISCs and newly formed EBs were stabilised and stem cell proliferation was prevented (Choi et al., 2011). Although the signals linking insulin signalling to junction disassembly and non-autonomous proliferation remain to be characterised, it is tempting to speculate that α-catenin recruitment of active vinculin might help strengthen adherens junctions when cells are under external tension, as would be the case during periods of active growth. In support of this hypothesis, upon refeeding (and thus active insulin signalling) after a week of starvation, *vinc*^*-/-*^ guts showed a recovery in size that initially surpassed the original one, correlating with increased cell production facilitated by junction disassembly. It will be interesting in future to establish whether in this system vinculin acts in parallel or downstream of the insulin signalling pathway.

Notably, E-cadherin mediated cell adhesion is also required in enterocytes to prevent stem cell renewal, but in this case, by preventing the secretion of mitogenic signals (Liang et al., 2017). Such coupling of differentiated cell density and self-renewal was also demonstrated *in vivo* in the mouse adult stratified epidermis, where stem cells divide in response to delamination (Mesa et al., 2018).

### Mechanosensing in intestinal precursors regulates tissue homeostasis

It seems fitting that proteins activated in response to tissue tension couple differentiation to self-renewal. Strikingly, we observed regional differences in the rate of tissue turnover in the posterior midgut of *vinc*^*102*.*1*^ guts. As the region with highest turnover in the mutant corresponded to a narrowing portion of the gut where the domains R4 and R5 are separated by a constriction of the intestinal wall (cartoon fig 1B, Buchon et al., 2013; Marianes & Spradling, 2013), we speculate that a local increase in mechanical forces (for example through visceral muscle contractions and food passage) induce vinculin mediated junction reinforcement to prevent untimely cell production. Consistent with this, β-catenin had a stronger cortical localisation in this area compared to more anterior areas of the midgut (data not shown). Another hypothesis is that this gut area presents with differential expression of signalling components. These issues remain to be solved.

Our work demonstrated that vinculin is required in EBs to prevent their precocious activation and differentiation into absorptive cells. How vinculin relates to transcription factors activated at this stage such as Esg and Klu remains to be formally tested. However, we observed that *vinc* knockdown in EBs cells led them to stall half-way through differentiation, whilst triggering proliferation in adjacent ISCs. This suggests that vinculin prevents the premature activation of dormant EBs. Once past a certain threshold of mechanical force, vinculin might not be sufficient to prevent tissue growth. Alternatively, vinculin might also contribute to terminal differentiation, either by direct signalling of terminal differentiation or by sensing local tissue density in a similar fashion as contact inhibition (Alegot et al, 2019; Biswas et al, 2021). Indeed, gut stretching after food ingestion induces proliferation through activation of the YAP/TAZ homolog Yorkie. The mechanical stress induces a displacement of the Ste20 kinase Misshapen from the membrane of EBs, thereby relieving interactions with upstream mediators of the Hippo pathway, and thus allowing Yorkie nuclear translocation, activation of target genes and midgut growth (Li et al., 2018). We suggest that vinculin contributes to mechanical force sensing in EBs and more generally helps coordinate self-renewal and differentiation into absorptive cells. Interestingly, the precursors of enteroendocrine cells can also transduce mechanical stress to control differentiation. In this case, however, cell fate is acquired through increased intracellular calcium downstream of the stretch-activated channel piezo expressed in EE precursors (He et al., 2018).

Thus, lineage commitment in the *Drosophila* intestine seems to depend on mechano-sensing pathways specifically active in precursors cells. Together with the role of integrin activation in intestinal stem cells to regulate self-renewal (this work; Lin et al., 2013), it becomes apparent cell-type specific mechano-sensing pathways contribute to fine tuning cell production and tissue composition.

### Outlook

In the mammalian intestine, stem cells are located in crypts whilst progenitors concentrate at the bottom of villi, which are populated by differentiated cells. Epithelial turnover is dependent on E-cadherin mediated adhesion (Hermiston & Gordon, 1995; Hermiston et al., 1999) and continuous cell migration (Krndija et al., 2019); thus, it is harder to establish if forces contribute to proliferation and lineage commitment or active cell migration (and indirectly self-renewal). The mechanoeffector protein vinculin is deregulated in gastrointestinal diseases, with expression levels reduced in colorectal cancers (Goldmann et al., 2013; Li et al., 2014) and auto anti-vinculin antibodies are produced in inflammatory bowel syndrome (Rezaie et al., 2017). The use of a simpler model organism, with a flat intestinal epithelium where stem cells are dispersed among their progeny (Micchelli & Perrimon, 2006; Ohlstein & Spradling, 2006) and cell migration is sporadic (Antonello et al., 2015; Martin et al., 2018) has allowed us to identify an EB specific role for vinculin mediated by α-catenin and independent of integrin adhesion. It will be important in future to develop tools to experimentally define how and when vinculin is activated to strengthen cell junctions in the healthy intestine and how to best harness this data with regards to vinculin deregulation in disease.

## Material and Methods

### Fly strains and experimental conditions

All *Drosophila* stocks were maintained at 18°C or amplified at 25°C on standard medium (cornmeal, yeast, glucose, agar, water, Nipagin food medium). The following strains were used: *yw, Vinc-GFP* (Klapholz et al., 2015), *Vinc-RFP* (Klapholz 2015), *vinc*^*102*.*1*^ (Klapholz et al., 2015; a deletion that removes *vinc, SmydA-8* and partially *Mct1*), UAS-Vinc^CO^-RFP (Maartens et al 2016), UAS-sqh^DD^, UAS-sqhRNAi, UAS-zipperRNAi (gifts from Silvia Aldaz), *esg*^*ts*^*-FlipOUT* (Jiang et al., 2009), *esg*^*ts*^*Gal4* (expressed in progenitor cells, Jiang et al, 2009), esg-Gal4^ts^Su(H)Gal80 (expressed in ISCs, Wang et al., 2014), esg^ts^-REDDM (Antonello et al., 2015), Klu^ts^-REDDM (expressed in enteroblasts, Antonello et al, 2015), MyoIA^ts^Gal4 (expressed in ECs, Jiang et al., 2009), *vinc* RNAi #25965 (Bloomington, TRiP.JF01985), *vinc* RNAi #41959 (Bloomington, TRiP.HMS02356), NRE-lacZ (Furriols and Bray, 2001), 10XSTAT-GFP (Bach et al., 2007). mys RNAi #27735 (Bloomington, TRIP.JF02819), talinRNAi #28950 (Bloomington, TRIP.HM05161), talin RNAi #33913 (Bloomington, TRIP.HMS00856), α-Cat RNAi #33430 (Bloomington, TRIP.HMS00317), α-Cat RNAi #38987 (Bloomington, TRIP.HMS00837). The following lines were gifted by K. Irvine (Alegot et al., 2019): α-Cat-FL-V5; α-Cat RNAi #33430, α-CatΔM1a-v5; α-Cat RNAi #33430, α-CatΔM1b-v5; α-Cat RNAi #33430, α-CatΔM2-v5; α-Cat RNAi #33430 - See Supplementary material for complete genotypes.

#### RNAi knockdown

RNAi knockdown efficiencies were tested by immunostaining (*mys, talin* and α*-Catenin*) or by RT-qPCR (*vinc* RNAis). Transcripts were downregulated by UAS-driven RNAi conditional expression in various cell populations using appropriate Gal4 driver lines combined with the thermosensitive repressor Gal80, to restrict expression to just adult stages, by shifting to the permissive temperature 29°C. To prevent RNAi expression during development, flies were kept at restrictive temperature (18°C) until 3 days after hatching, after which 10-15 mated females were transferred at the permissive temperature for a minimum of 14 days to allow intestinal expression of UAS-driven constructs. Food was changed every 2 days throughout the experiment.

#### Escargot-Flip-Out experiments (fig. 2C-G)

*flippase* expression under the control of *esg-Gal4* in progenitor cells, at the permissive temperature of the repressor gal80 (29°C), excises a transcriptional stop between a ubiquitous *actin* promotor and Gal4, resulting in permanent, heritable expression of Gal4 in all progenitor cells and their progeny, as revealed with UAS-GFP (Jiang et al, 2009, fig.2C and suppl fig.3A). Flies were kept at restrictive temperature (18°C) until 3 days after hatching, after which 10-15 males were transferred at the permissive temperature for a minimum of 14 days. Food was changed every 2 days throughout the experiment.

#### ReDDM experiments (figs. 4, 5)

The lineage tracer ReDDM (<u>R</u>epressible <u>D</u>ual <u>D</u>ifferential stability cell <u>M</u>arker) is a UAS-driven transgene encoding a mCD8-GFP with a short half-life, and H2B-RFP with a long half-life both contained in two independent UAS-constructs (Antonello et al., 2015). When expressed under the control of *esg-Gal4* or *Klu-Gal4*, this system allowed us to identify unambiguously newly differentiated enterocytes as RFP^+^GFP^-^ cells and thus establish the rate of differentiation.

### Dissection and Immunostaining

Adult guts were dissected in PBS and fixed twice during 20 min in fresh 4% paraformaldehyde diluted in PBS (18814-10, Polysciences) with a 10 min wash in PBT 0,1% (PBS + Triton X100 Sigma Aldrich) in between, apart for fig.2B-F, where guts were fixed once. Samples were then washed 5 min three times in PBT 0.1% and permeabilised for 30 min in PBT 1%. All samples were then incubated 30min at room temperature in blocking buffer (PBS + 1% BSA, A2153-10G, Sigma) followed with primary antibodies incubation in blocking buffer overnight at 4°C. Samples were then washed 3 times 15min in PBT 0.1%, subjected to secondary antibody staining in blocking buffer for 2 hours at room temperature followed by 3 washings in PBT 0.1%. Guts were mounted in Vectashield (H-100, Vector Laboratories) between coverslip and glass slide. The following primary antibodies were used: chicken anti-GFP, 1/1000 (Abcam 13970); mouse anti-Delta, 1/1000 (C594.9B-c, DSHB); chicken anti-Beta galactosidase, 1/1000 (Abcam 9361); mouse anti-Armadillo (β-Catenin), 1/50 (N2 7A1-c, DSHB), mouse anti-V5, 1/400 (R96025, ThermoFisher), rat anti-α--Catenin, 1/20 (DSHB D-CAT1), rabbit anti-PH3(Ser10), 1/500 (9701 CST); mouse anti-Prospero, 1/50 (DSHB, MR1a). Alexa-488, 555 and 647 conjugated secondary goat antibodies or Phalloidin were used (Molecular probes) and nuclei were counterstained with DAPI.

### Imaging and analysis

All confocal images were taken on a Leica SP8 Confocal microscope with x20 air lens, x40 and x63 oil immersion lenses. 405nm, 488nm, 516nm and 633nm lasers were used as appropriate, with gain and power consistent within experiments. For z-stacks, images were acquired every 1μm through the tissue. Whole gut photographs were taken on a Zeiss Stemi 305 connected to an iPad or a Leica M165FC microscope with Leica DFC3000G camera. All images were analysed with FiJi and Adobe Photoshop CS. When necessary, images were stitched using the ImageJ pairwise stitching plugin.

### Feeding, starvation, refeeding regime and survival

*yw* and *vinc*^*102*.*1*^ stocks were reared in parallel in bottles of standard medium at room temperature for several generations prior to the experiment. Flies used in the experiment underwent embryonic, larval and pupal development in bottles of standard medium at room temperature, except for the final two days before hatching when bottles were placed at 25°C to speed up development. After discarding early emergents, bottles were placed back at 25°C for two days, after which all synchronised hatchlings were collected and transferred to new bottles of standard medium. These bottles were placed at 25°C under a 12 hour light 12 hour dark cycle for two days to allow flies to reach sexual maturity and mate. Flies were then anesthetised for minimal time to retrieve female flies, which were transferred into vials of standard medium at a density of 12 flies per vial. Vials were placed horizontally at 25°C under a 12 hour light 12 hour dark cycle, with vial position randomised to control for any variation in temperature or humidity in the incubator. Survival was scored each morning, and flies were transferred to new vials containing fresh food every two days. When flies were 7 days old, 12 flies per genotype were dissected in PBS (BR0014G, Oxoid) and imaged immediately to measure midgut length (between the centre of the proventriculus and the Malpighian tubule attachment site, between arrows in fig. 6A) and width in the R4/5 region. Of the remaining flies for each genotype, approximately half were transferred to starvation conditions – vials containing a cellulose acetate bung (Fly1192, Scientific Laboratory Supplies) soaked in mineral water (Shropshire Hills Mineral Water, Wenlock Spring Water Limited). The bung was replaced every 24 hours to prevent drying. Control flies were continuously fed in vials of standard medium. After 7 days of starvation, 12 flies from each genotype and condition (‘fed-starved’ and ‘continuously fed’) were dissected and midgut length and width measured. Remaining ‘fed-starved’ flies were then transferred back into vials of standard medium. After four days of refeeding, 12 flies from each genotype and condition (‘fed-starved-refed’ and ‘continuously fed’) were dissected and midgut length and width measured. All remaining flies continued to be reared in vials of standard medium with dissections performed every three-four days, until the flies reached 28 days old at which point the experiment ended. Throughout the starvation and refeeding assay, survival was scored each morning. Any flies which left the experiment prior to natural death (e.g. due to escape or squashing) were censored and removed from the analysis. The survival graph was constructed assuming that dissected flies would have showed the same percentage survival as remaining flies, had they been allowed to continue living.

### Quantifications and Statistics

Unless otherwise indicated the images shown in this study and the respective quantifications focused on the specific R4/5 region of the posterior midgut. ISC, EB and progenitor ratios were quantified manually through the cell counter plugin in ImageJ software and represented as the ratio of GFP^+^ cells over the total number of nuclei in DAPI. Cell density was assessed semi automatically through binarization of DAPI signal in ImageJ and measurement of the number of nuclei in a fixed area (1000 µm^2^) representative of the whole images processed. The differentiation index was assessed by manually counting GFP^+^RFP^+^ and GFP^-^RFP^+^ cells via the cell counter plugin in ImageJ. The differentiation index was calculated as the ratio of differentiated cells (GFP^-^RFP^+^) number over progenitors (GFP^+^RFP^+^) number. Nuclear size was measured by binarizing the DAPI signal of all nuclei using ImageJ. Cell type was defined using anti-Dl (ISC) antibody staining, Su(H)-lacZ (EB) reporter. To exclude enteroendocrine cells from EC nuclear size measurement only Dl^-^, Su(H)^-^ and bigger than ISC nuclear size were considered. For cross-sectional nuclei counting, confocal z-stacks through the full midgut depth were visualised with the ImageJ orthogonal view tool. Number of epithelial cell nuclei and maximum midgut width were measured in progressive cross-sections every 30μm along 300μm of the R4/5 region for both genotypes. Data were plotted with Graphpad Prism 7, 8 and 9 software. In all graphs, results represent median values, error bars represent interquartile range. Statistical significance was calculated using a Mann-Whitney test and p values <0.05 were considered statistically significant. Survival was compared between *yw* and *vinc*^*102*.*1*^ flies using the Logrank test.

### Wing mounting

Wings were mounted in Euparal (R1344A, Agar Scientific). on a glass slide (SuperFrost 1.0mm, ISO 8037/1, VWR International) with coverslip (22×50mm#1, 12342118, Menzel-Gläser). Images were taken on a Zeiss Axiophot microscope with a Q Imaging QICAM FAST1394 camera.

### Scanning electron microscopy

Guts of 7-day old female flies were dissected in Schneider’s medium (S3652, Sigma) and fixed overnight at 4°C in 2% glutaraldehyde, 2% formaldehyde in 0.05M sodium cacodylate buffer (pH7.4) containing 2 mM CaCl_2_. After fixation, guts were cut at the middle point of the posterior midgut with a razor blade (WS1010, Wilkinson Sword). Guts were washed with 0.05M sodium cacodylate buffer and osmicated overnight at 4°C in 1% OsO_4_, 1.5% potassium ferricyanide, 0.05M sodium cacodylate buffer (pH 7.4). Guts were washed in deionised water (DIW) and treated for 20minutes in the dark at room temperature with 0.1 % thiocarbohydrazide/DIW. Guts were washed again in DIW and osmicated for 1 hour at room temperature in 2% OsO_4_ in DIW. Following washing in DIW, guts were treated for 3 days at 4°C in 2% uranyl acetate in 0.05M maleate buffer (pH 5.5). Guts were washed in DIW and stained for one hour at 60°C in lead aspartate solution (0.33 g lead nitrate in 50ml 0.03M aspartic acid solution (pH 5.5)). Following washing in DIW, guts were dehydrated in 50, 70, 95 & 100% ethanol, 3 times in each for at least 5 minutes each. Guts were then dehydrated twice in 100% dry ethanol, twice in 100% dry acetone and three times in dry acetonitrile for at least 5 minutes each. Guts were placed for 2 hours at room temperature in a 50/50 mixture of Quetol (TAAB) resin mix and 100% dry acetonitrile. Guts were incubated in pure Quetol resin mix (12 g Quetol 651, 15.7 g NSA, 5.7 g MNA and 0.5 g BDMA (all from TAAB)) for 5 days, replacing with fresh resin mix each day. Guts were embedded in coffin mounds and incubated for 48 hours at 60°C. Resin-embedded samples were mounted on aluminium SEM stubs using conductive epoxy resin and sputter-coated with 35nm gold. Blockfaces were sectioned using a Leica Ultracut E and coated with 30nm carbon for conductivity. The samples were imaged in a FEI Verios 460 SEM at an accelerating voltage of 3-4 keV and a probe current of 0.2 pA in backscatter mode using the CBS-detector in immersion mode. Large area maps were acquired using FEI MAPS software for automated image acquisition. MAPS settings for high-resolution maps were 1536×1024 pixel resolution, 10μs dwell time, 2 line integrations, magnification ∼8000x, working distance ∼ 4mm, tile size 15.9μm, default stitching profiles.

### Fluorescence heatmaps

Posterior midguts were imaged with a Leica M165FC microscope with Leica DFC3000G camera. Fluorescence intensity was measured along three lines of defined position (‘upper quarter’, ‘middle’ and ‘lower quarter’) spanning the length of the posterior-most 1200μm of each midgut using the ImageJ plot profile tool. Fluorescence intensity along the three lines was averaged for each gut. Then average fluorescence intensity was calculated for each 50μm gut segment and normalised by dividing by the lowest fluorescence value (to account for variation in image brightness within and between genotypes). This was repeated for tens of guts for each genotype, and the overall average normalised values for each 50μm gut segment were plotted as a heatmap using Prism.

## Acknowledgements

We thank Nick Brown, Sarah Bray, Tobias Reiff, Maria Dominguez, Cedric Polesello and Ken Irvine for kindly sharing flies with us. *Drosophila* stocks used in this study were otherwise obtained from the Bloomington *Drosophila* Stock Center (NIH P40OD018537). Several antibodies were obtained from the Developmental Studies Hybridoma Bank, created by the NICHD of the NIH and maintained at The University of Iowa, Department of Biology, Iowa City, IA 52242. We thank the Cambridge Advanced Imaging Centre for their help with electron microscopy and for use of the confocal facility. We are grateful to members of the Brown lab (PDN, Cambridge) for insightful suggestions and comments on the project and the manuscript, and Aki Stubb (PDN, Cambridge) for constructive feedback. This work was supported by a Sir Henry Dale Fellowship jointly funded by the Wellcome Trust and the Royal Society to G.K. [Grant number 206208/Z/17/Z] and a Wellcome Trust PhD studentship to B.L.E.T. [Programme number 102175/B/13/Z].

## Author contributions

JB, BLET and GK conceptualised, conducted experiments and analysed the data. JB and BLET prepared figures. GK wrote the manuscript, with editing contributions from JB and BLET. GK supervised and acquired funding. All authors approved the submitted article.

## Open access

For the purpose of open access, the author has applied a CC BY public copyright licence to any Author Accepted Manuscript version arising from this submission.

## Supplemental information

### Supplemental Genotypes

**Figure 1:**

B-F:

<u>Ctrl:</u> y,w / w; esgGal4, tubGal80[ts] UAS GFP / + ; + / +

<u>mysRNAi #27735:</u> y,w / y,v ; esgGal4, tubGal80[ts] UAS GFP / + ; P{y[+t7.7] v[+t1.8]=TRiP.JF02819}attP2 / +

<u>talinRNAi #28950:</u> y,w / y,v ; esgGal4, tubGal80[ts] UAS GFP / P{y[+t7.7] v[+t1.8]=TRiP.HM05161}attP2 ; + / +

<u>talinRNAi #33913:</u> y,w / y,sc,v,sev ; esgGal4, tubGal80[ts] UAS GFP / P{y[+t7.7] v[+t1.8]=TRiP.HMS00856}attP2 ; + / +

<u>vincRNAi #25965:</u> y,w / y,v ; esgGal4, tubGal80[ts] UAS GFP / + ; P{y[+t7.7] v[+t1.8]=TRiP.JF01985}attP2 / +

<u>vincRNAi #41959:</u> y,w / y,sc,v,sev ; esgGal4, tubGal80[ts] UAS GFP / P{y[+t7.7] v[+t1.8]=TRiP.HMS02356}attP2 ; + / +

**Figure 2:**

A-B:

<u>yw:</u> yw / yw ; + / + ; + / +

<u>vinc^102.1^:</u> vinculin[102.1]/ vinculin[102.1] ; + / + ; + / +

D-G:

<u>yw esg^ts^ F/O:</u> yw/ Y ; esg-Gal4, tubGal80[ts], UAS-GFP / + ; UAS-flp, act>CD2>Gal4 / +

<u>vinc^102.1^ esg^ts^ F/O:</u> vinculin[102.1] / Y ; esg-Gal4, tubGal80[ts], UAS-GFP / + ; UAS-flp, act>CD2>Gal4 / +

<u>vinc^102.1^, vincRFP esg^ts^ F/O:</u> vinculin[102.1] / Y ; esg-Gal4, tubGal80[ts], UAS-GFP / vincRFP ; UAS-flp, act>CD2>Gal4 / +

**Figure 3:**

A-C:

<u>Ctrl:</u> w / yw ; esg-Gal4 UAS-YFP / + ; Su(H) GBE-Gal80, tubGal80ts / +

<u>vincRNAi #25965:</u> y,v / w ; esg-Gal4 UAS-YFP / + ; P{y[+t7.7] v[+t1.8]=TRiP.JF01985}attP2 / Su(H) GBE-Gal80, tubGal80ts

<u>mysRNAi #27735:</u> y,v / w ; esg-Gal4 UAS-YFP / + ; P{y[+t7.7] v[+t1.8]=TRiP.JF02819}attP2 / Su(H) GBE-Gal80, tubGal80ts

<u>talinRNAi #28950:</u> y,v / w ; esg-Gal4 UAS-YFP / P{y[+t7.7] v[+t1.8]=TRiP.HM05161}attP2 ; Su(H) GBE-Gal80, tubGal80ts / + D-F:

<u>Ctrl:</u> hsflp, FRT19A, tubGal80 / FRT19A ; UAS-mCD8-GFP / + ; tub-Gal4 / +

<u>Vinc^102.1^:</u> hsflp, FRT19A, tubGal80 / FRT19A vinc^102.1^ ; UAS-mCD8-GFP / + ; tub-Gal4 / +

<u>vincRNAi #25965:</u> hsflp, FRT19A, tubGal80 / FRT19A ; UAS-mCD8-GFP / + ; tub-Gal4 / P{y[+t7.7] v[+t1.8]=TRiP.JF01985}attP2

**Figure 4:**

A-D:

<u>yw, NRE-lacZ:</u> yw / Y ; NRE-lacZ [3-45] / + ; + / +

<u>vinc^102.1^, NRE-lacZ:</u> vinculin[102.1]/ Y ; NRE-lacZ [3-45] / + ; + / + F,G:

<u>Ctrl:</u> yw / w ; esg-Gal4, UAS-mCD8-GFP / + ; tubGal80[ts], UAS H2B RFP / +

<u>vincRNAi #25965:</u> yv / + ; esg-Gal4, UAS-mCD8-GFP / + ; tubGal80[ts], UAS H2B RFP / P{y[+t7.7] v[+t1.8]=TRiP.JF01985}attP2 I,J:

<u>Ctrl:</u> yw / w; UAS-CD8-GFP, tubGal80[ts] ; Klu-Gal4, UAS-H2B-RFP / SM6a-TM6B

<u>vincRNAi #25965:</u> yv / w ; UAS-CD8-GFP, tubGal80[ts] / + ; Klu-Gal4, UAS-H2B-RFP / P{y[+t7.7] v[+t1.8]=TRiP.JF01985}attP2

**Figure 5:**

B,C:

<u>Ctrl:</u> y,w / w ; esgGal4, tubGal80[ts] UAS GFP / + ; + / +

<u>αcatRNAi #33430:</u> y,sc,v,sev / w ; esgGal4, tubGal80[ts] UAS GFP / + ; P{y[+t7.7] v[+t1.8]=TRiP.HMS00317}attP2 / +

<u>αcatRNAi #38987:</u> y,v / w ; esgGal4, tubGal80[ts] UAS GFP / P{y[+t7.7] v[+t1.8]=TRiP.HMS01903}attP40 ; + / +

<u>vincRNAi #25965:</u> y,w / y,v ; esgGal4, tubGal80[ts] UAS GFP / + ; P{y[+t7.7] v[+t1.8]=TRiP.JF01985}attP2 / +

E-G:

<u>Ctrl:</u> yw / w ; esg-Gal4, UAS-mCD8-GFP / + ; tubGal80[ts], UAS H2B RFP / +

<u>αcatFL + αcatRNAi #33430:</u> w / + ; esg-Gal4, UAS-mCD8-GFP / UAST-αCatFLmut-V5[attP40] ; tubGal80[ts], UAS H2B RFP / P{y[+t7.7] v[+t1.8]=TRiP.HMS00317}attP2

<u>αcatΔM1a + αcatRNAi #33430:</u> w / + ; esg-Gal4, UAS-mCD8-GFP / UAST-αCatΔ265-397mut-V5[attP40] ; tubGal80[ts], UAS H2B RFP / P{y[+t7.7] v[+t1.8]=TRiP.HMS00317}attP2

<u>αcatΔM1b + αcatRNAi #33430:</u> w / + ; esg-Gal4, UAS-mCD8-GFP / UAST-αCatΔ273-398mut-V5[attP40] ; tubGal80[ts], UAS H2B RFP / P{y[+t7.7] v[+t1.8]=TRiP.HMS00317}attP2

<u>αcatΔM2 + αcatRNAi #33430:</u> w / + ; esg-Gal4, UAS-mCD8-GFP / UAST-αCatΔ397-509mut-V5[attP40] ; tubGal80[ts], UAS H2B RFP / P{y[+t7.7] v[+t1.8]=TRiP.HMS00317}attP2

**Figure 6:**

A-D:

<u>yw:</u> yw / yw ; + / + ; + / +

<u>vinc^102.1^:</u> vinculin[102.1]/ vinculin[102.1] ; + / + ; + / +

**Supplementary figure 1:**

<u>Vinc-GFP:</u> +/+; vinc-GFP/vinc-GFP; +/+; +/+

**Supplementary figure 2:**

<u>yw:</u> yw / yw ; + / + ; + / +

<u>vinc^102.1^:</u> vinculin[102.1]/ vinculin[102.1] ; + / + ; + / +

**Supplementary figure 3:**

B: <u>vinc^102.1^ esg^ts^ F/O:</u> vinculin[102.1] / Y ; esg-Gal4, tubGal80[ts], UAS-GFP / + ; UAS-flp, act>CD2>Gal4 / +

**Supplementary figure 4:**

<u>Ctrl:</u> hsflp, FRT19A, tubGal80 / FRT19A ; UAS-mCD8-GFP / + ; tub-Gal4 / +

<u>Vinc^102.1^:</u> hsflp, FRT19A, tubGal80 / FRT19A vinc^102.1^ ; UAS-mCD8-GFP / + ; tub-Gal4 / +

<u>vincRNAi #25965:</u> hsflp, FRT19A, tubGal80 / FRT19A ; UAS-mCD8-GFP / + ; tub-Gal4 / P{y[+t7.7] v[+t1.8]=TRiP.JF01985}attP2

**Supplementary figure 5:**

<u>yw, 10XSTAT-GFP</u>: w / Y ; 10XSTAT-GFP ; + / +

<u>vinc^102.1^, 10XSTAT-GFP:</u> vinculin[102.1] / Y ; 10XSTAT-GFP ; + / +

**Supplementary figure 6:**

<u>Ctrl:</u> yw / w; UAS-CD8-GFP, tubGal80[ts] ; Klu-Gal4, UAS-H2B-RFP / SM6a-TM6B

<u>vincRNAi #25965:</u> yv / w ; UAS-CD8-GFP, tubGal80[ts] / + ; Klu-Gal4, UAS-H2B-RFP / P{y[+t7.7] v[+t1.8]=TRiP.JF01985}attP2

**Supplementary figure 7:**

<u>Ctrl:</u> yw/ yw ; MyoIA-Gal4 tub-Gal80ts / UAS-CD8-GFP ; + / +

<u>vincRNAi #25965:</u> yw/ yv ; MyoIA-Gal4 tub-Gal80ts / UAS-CD8-GFP ; P{y[+t7.7] v[+t1.8]=TRiP.JF01985}attP2 / +

**Supplementary figure 8:**

hsflp, FRT19A, tubGal80 / FRT19A ; UAS-mCD8-GFP / + ; tub-Gal4 / P{y[+t7.7] v[+t1.8]=TRiP.JF01985}attP2

**Supplementary figure 9:**

<u>yw esg^ts^ F/O:</u> yw/ Y ; esg-Gal4, tubGal80[ts], UAS-GFP / + ; UAS-flp, act>CD2>Gal4 / +

<u>yw esg^ts^ F/O + UASVinc^CO^-RFP:</u> yw/ Y ; esg-Gal4, tubGal80[ts], UAS-GFP / UAS-Vinc^CO^-RFP ; UAS-flp, act>CD2>Gal4 / +

<u>vinc102.1 esg^ts^ F/O:</u> vinculin[102.1] / Y ; esg-Gal4, tubGal80[ts], UAS-GFP / + ; UAS-flp, act>CD2>Gal4 / +

<u>vinc102.1 esg^ts^ F/O + UASVinc^CO^-RFP:</u> vinculin[102.1] / Y ; esg-Gal4, tubGal80[ts], UAS-GFP / UAS-Vinc^CO^-RFP ; UAS-flp, act>CD2>Gal4 / +

**Supplementary figure 10:**

<u>Ctrl:</u> yw / w ; esg-Gal4, UAS-mCD8-GFP / + ; tubGal80[ts], UAS H2B RFP / +

<u>sqh^DD^:</u> w / + ; esg-Gal4, UAS-mCD8-GFP / UAS-sqh^DD^ ; tubGal80[ts], UAS H2B RFP / +

<u>sqh-RNAi:</u> w / + ; esg-Gal4, UAS-mCD8-GFP / UAS-sqhRNAi ; tubGal80[ts], UAS H2B RFP / +

<u>zipper-RNAi:</u> w / hs-flp ; esg-Gal4, UAS-mCD8-GFP /UAS-zipperRNAi ; tubGal80[ts], UAS H2B RFP / +

**Supplementary figure 11:**

<u>yw:</u> yw / yw ; + / + ; + / +

<u>vinc^102.1^:</u> vinculin[102.1]/ vinculin[102.1] ; + / + ; + / +

**Supplementary Figure 1:**
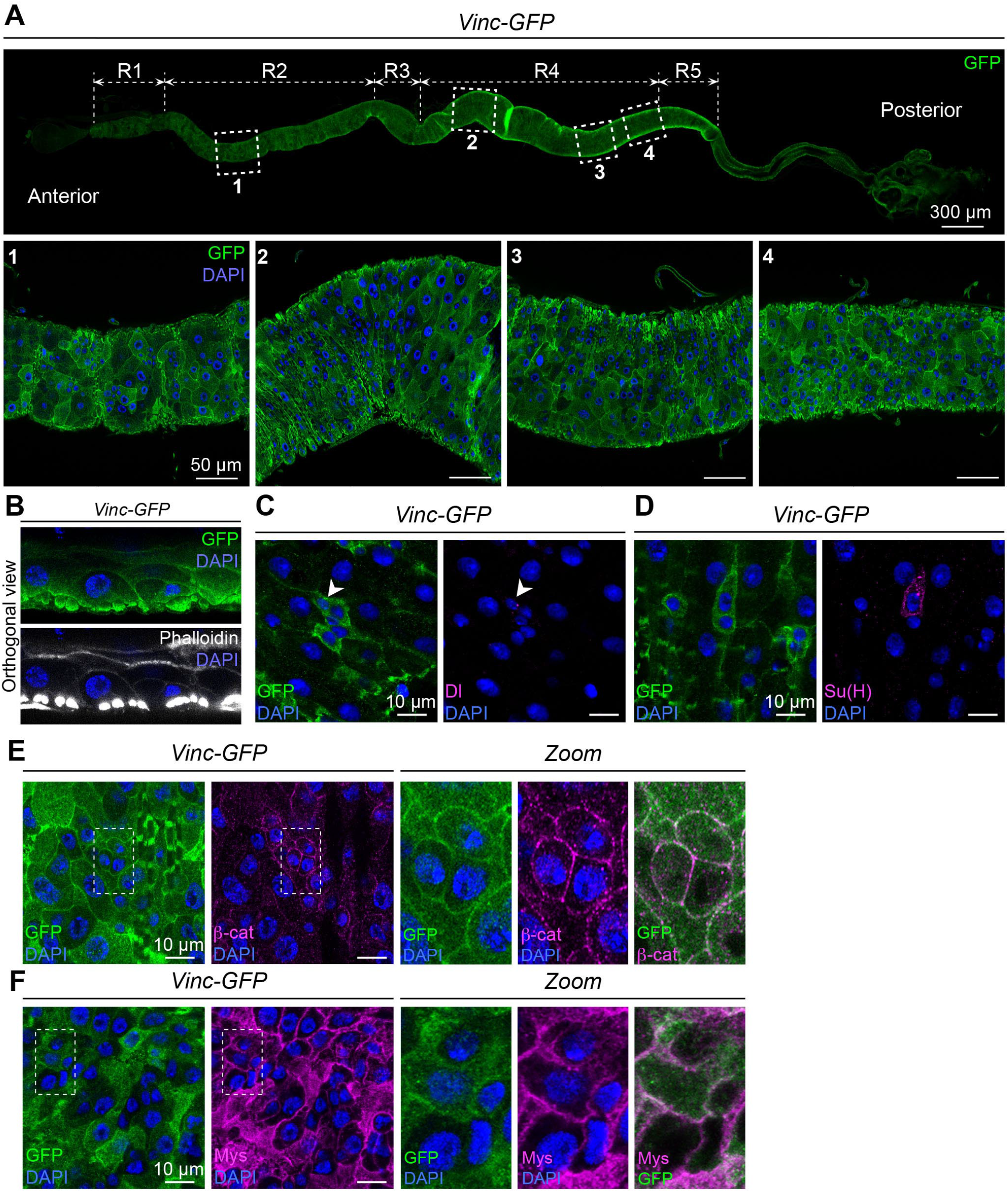
**(A)** Vinculin localisation along the whole gut (top panel). Vinculin tagged with GFP (*Vinc-GFP*, green) is encoded by a genomic construct. Bottom panels: close-ups in regions R2 (1), and R4 (2-4) of the intestine showing vinculin localisation in the epithelium. Nuclei are marked with DAPI (blue) throughout the figure. **(B)** Orthogonal view of the intestinal epithelium showing vinculin (Vinc-GFP, green) in epithelial cell membranes and in visceral muscles marked with Phalloidin (bottom panel, white). **(C)** Vinculin is expressed in ISCs marked with Delta (Dl, magenta). **(D)** Vinculin is expressed in EBs marked by expression of Su(H) (magenta). **(E)** Vinculin colocalizes with β-catenin (β-cat, magenta). Right panels present close-ups of individual and merged vinculin and β-catenin protein localization from dashed line area highlighted in left panel. **(F)** Vinculin colocalizes with integrin βPS subunit (Mys, magenta). Right panels present close-ups of individual and merged vinculin and Mys protein localization from dashed line area highlighted in left panel.

**Supplementary Figure 2:**
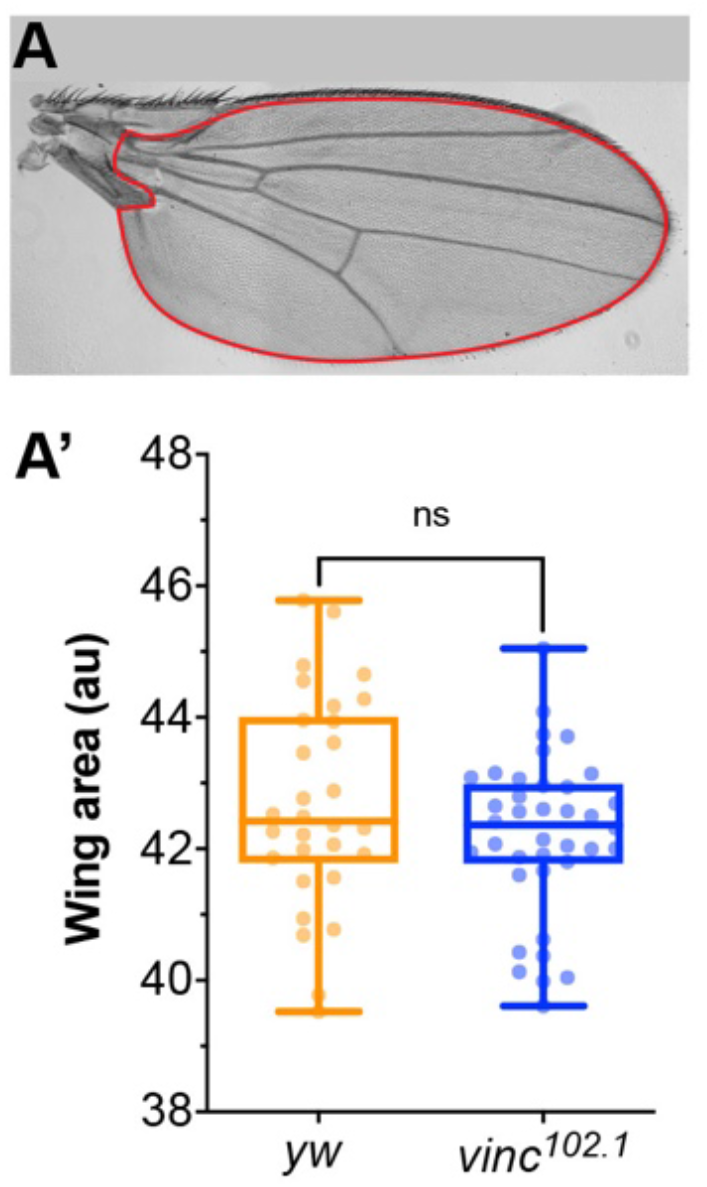
**(A)** Adult female wing from a *yw* fly shown as an example. Only the red area depicted was considered for size quantifications shown in A’. **(A’)** Quantification of wing area in *yw* and *vinc*^*102*.*1*^ female flies. *yw*: n=30 wings; *vinc*^*102*.*1*^: n=37 wings. Two-tailed Mann-Whitney test. ns: not significant.

**Supplementary Figure 3:**
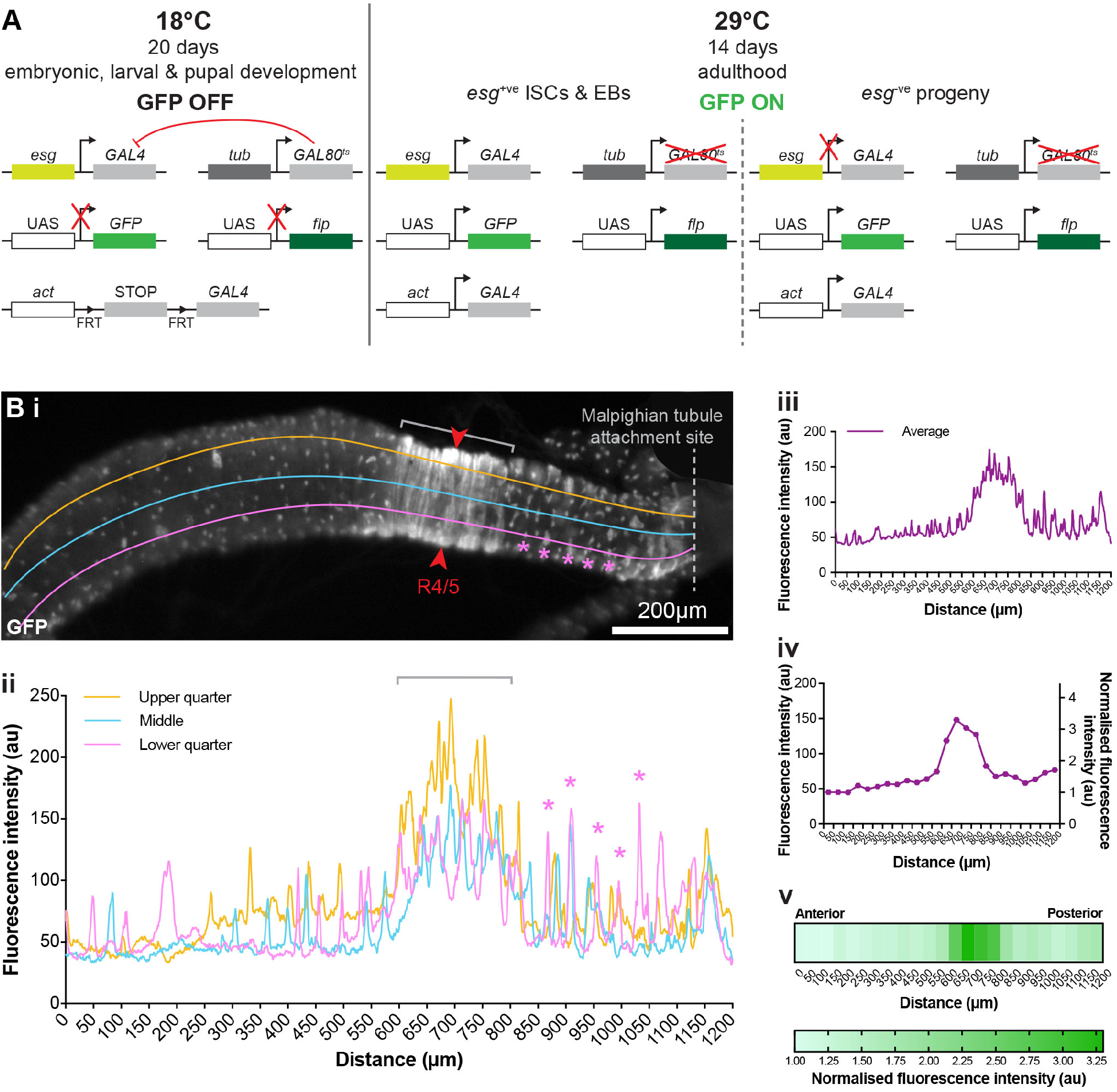
**(A)** Schematic of the genetic *esg*^*ts*^ *F/O* system used in this study. Flies were reared at non-permissive temperature (18°C) from embryo up until adult hatching. Therefore, during all developmental stages, the thermosentitive Gal4 inhibitor Gal80, expressed under the *tubulin* (*tub*) promoter, prevents GFP and flippase expression in *esg*^*+*^ cells. Three days old adult flies were then incubated at 29°C and this for 14 consecutive days. At this temperature the Gal80 is inactivated, allowing the concomitant expression of the GFP and the flippase in all *esg*^*+*^ cells. In the meantime, the STOP codon surrounded by two FRT cassettes is then excised activating constitutive expression of GFP in *esg*^*+*^ cells and their progeny under *actin* (*act*) promoter. **(Bi-Bv)** Workflow for generating fluorescence intensity heatmaps. **(i)** Example of a *vinc*^*102*.*1*^ *esg*^*ts*^ F/O gut showing GFP fluorescence intensity measurement at three positions, ‘upper quarter’ (orange line), ‘middle’ (blue line) and ‘lower quarter’ (pink line), along the posterior-most 1200 μm of each midgut. Red arrows indicate R4c and R4/5 regions of the posterior midgut where gut width was measured **(ii)** Example of a GFP intensity plot where individual GFP-positive clones appear as isolated peaks (pink asterisks) and the stereotypic extended GFP area, corresponding to increased intestinal cell renewal, appear as large continuous peaks (grey bracket) in *vinc*^*102*.*1*^ gut. **(iii)** Average of fluorescence intensity along the three lines (upper, middle and lower quarter) at each position. **(iv)** Average fluorescence intensity calculated for each 50μm gut segment and normalised by dividing by the lowest fluorescence value (to account for variation in image brightness within and between genotypes). **(v)** Normalised values plotted as a heatmap showing GFP intensity.

**Supplementary Figure 4:**
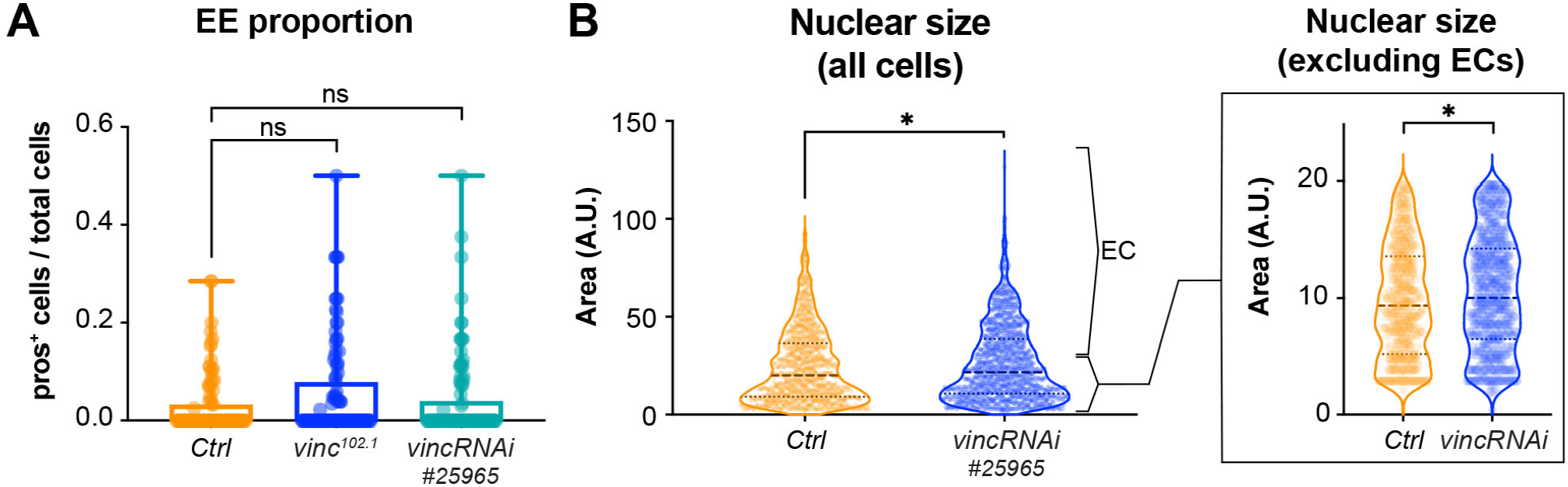
**(A)** Quantification of enteroendocrine cells (EEs) proportion in multicellular clones (number of pros^+^ cells/number of cells per clone). *Ctrl*: n=95 clones across 26 guts; *vinc*^*102*.*1*^: n=106 clones across 34 guts; *vincRNAi*^*#25965*^: n=102 clones across 25 guts. **(B)** Left plot: Quantification of clone cells nuclear size (DAPI area per cells). *Ctrl*: n=936 cells across 26 guts; *vincRNAi*^*#25965*^: n=1235 cells across 25 guts. Right plot: Same quantification of clone cells nuclear size shown in the left plot but excluding expected EC nuclear size data showing that ISC/EB pool nuclear size is still significantly different between *Ctrl* and *vincRNAi*^*#25965*^ clones. In A and B two-tailed Mann-Whitney test. ns: not significant, *: p<0.05.

**Supplementary Figure 5:**
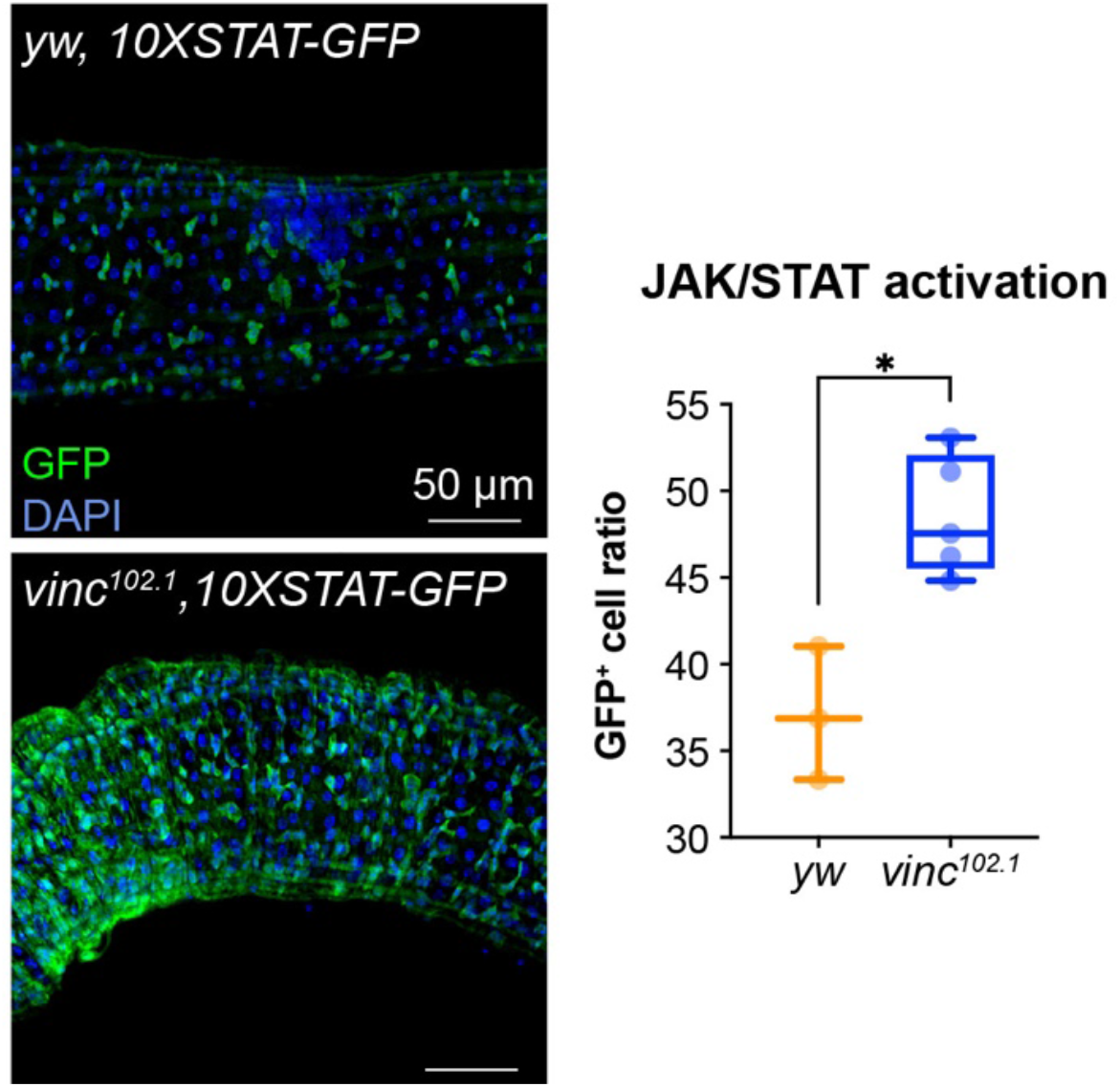
JAK-STAT reporter (10XSTAT-GFP) detected by GFP antibody staining (green) in 14-day-old *yw* or *vinc*^*102*.*1*^ guts of male flies. Nuclei are stained with DAPI (blue). Right graph present quantification of JAK/STAT activation (number of GFP^+^ cells/total number of cells) in *yw* and *vinc*^*102*.*1*^. Two-tailed Mann-Whitney tests were used for analysis. *: p<0.05.

**Supplementary Figure 6:**
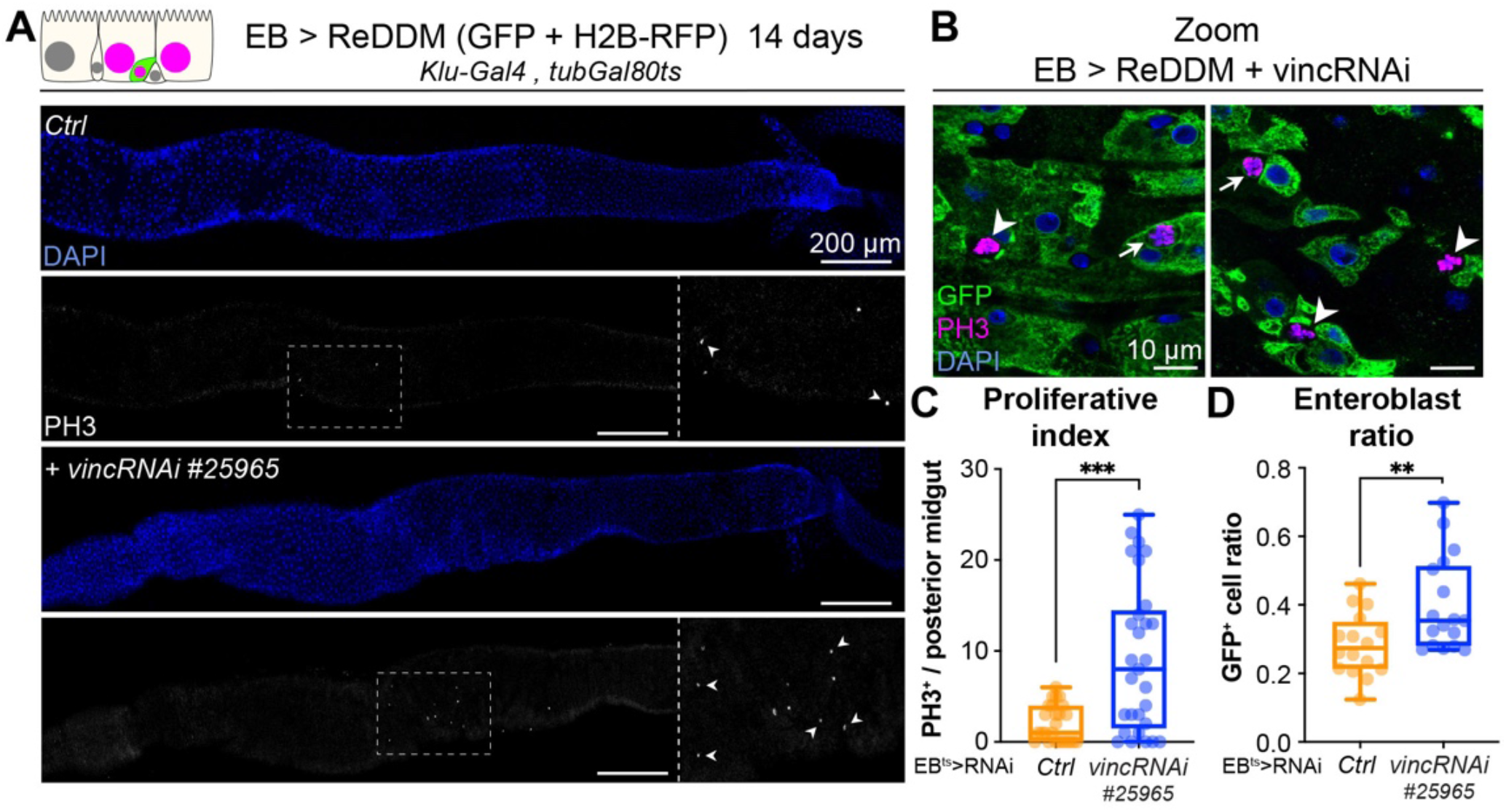
**(A)** Expression of ReDDM in EBs with *vincRNAi*^*#25965*^ or without (*Ctrl*) for 14 days. Note the increased number of mitotic cells (PH3^+^) when *vincRNAi*^*#25965*^ is expressed. Close ups show dashed line rectangles region with increased PH3^+^ cells (arrowheads) in *vincRNAi*^*#25965*^ expressing guts compare to *Ctrl*. Nuclei are stained with DAPI (blue). **(B)** Expression of ReDDM in EBs with *vincRNAi*^*#25965*^ for 14 days. Arrowheads indicate GFP^-^PH3^+^ dividing stem cells normally present in control conditions while arrows point at GFP^+^PH3^+^ dividing EB which can be observed after *vincRNAi*^*#25965*^ expression and not in control. Only GFP expression from the ReDDM system is shown here. Nuclei are stained with DAPI (blue). **(C)** Quantification of proliferative index (number of PH3^+^/posterior midgut) showing the increased proliferation following *vincRNAi*^*#25965*^ expression in EBs. *Ctrl*: n=25 guts; *vincRNAi*^*#25965*^: n=29 guts. **(D)** Quantification of the ratio of GFP^+^ EBs to the total number of cells after 14 days of ReDDM expression with *vincRNAi*^*#25965*^ or without (*Ctrl*). *Ctrl*: n=16 guts; *vincRNAi*^*#25965*^: n=17 guts. Mann-Whitney test, **: p<0.01.

**Supplementary Figure 7:**
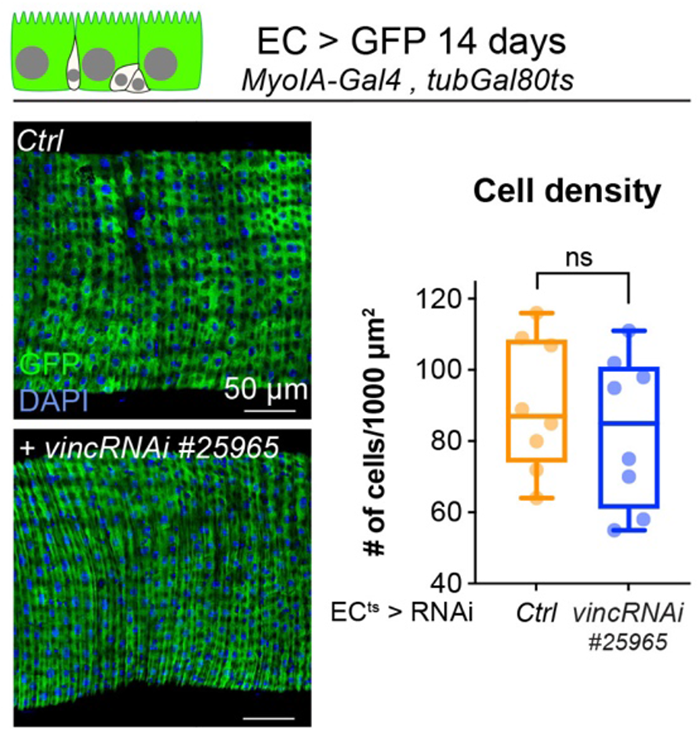
Effect of 14-day *vincRNAi*^*#25965*^ expression in ECs (GFP^+^) under the control of a thermosensitive Gal4 driver (*MyoIA-Gal4*^*ts*^) visualised in midguts stained for GFP (green) and DAPI (blue). Right graph represents the quantification of cell density (number of nuclei per 1000 μm^2^ area) from (E). *Ctrl*: n=8 guts; *vincRNAi*^*#25965*^: n=8 guts. Two-tailed Mann-Whitney tests were used for analysis. ns: not significant

**Supplementary Figure 8:**
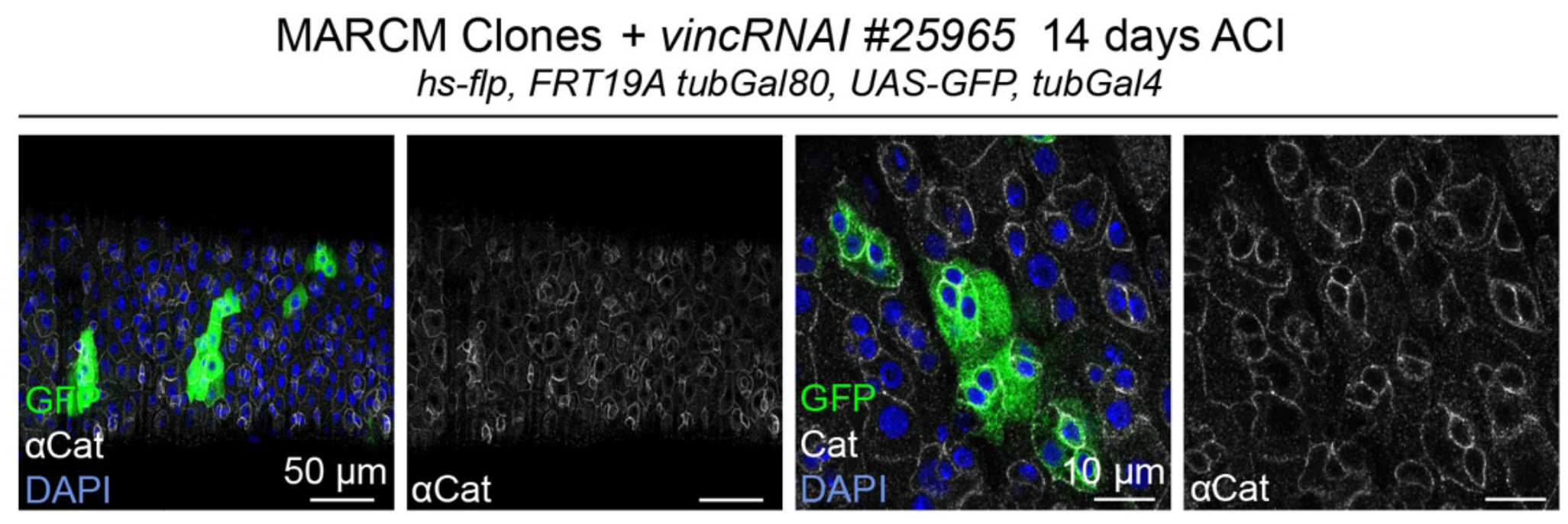
Examples of guts containing GFP-labelled mitotic clones *vinc*^*102*.*1*^ homozygous (GFP, green) stained for α-catenin (α-Cat, white). The distribution of a-Cat is indistinguishable between *vinc*^*102*.*1*^ clones and unlabelled control cells. Nuclei are stained with DAPI (blue).

**Supplementary Figure 9:**
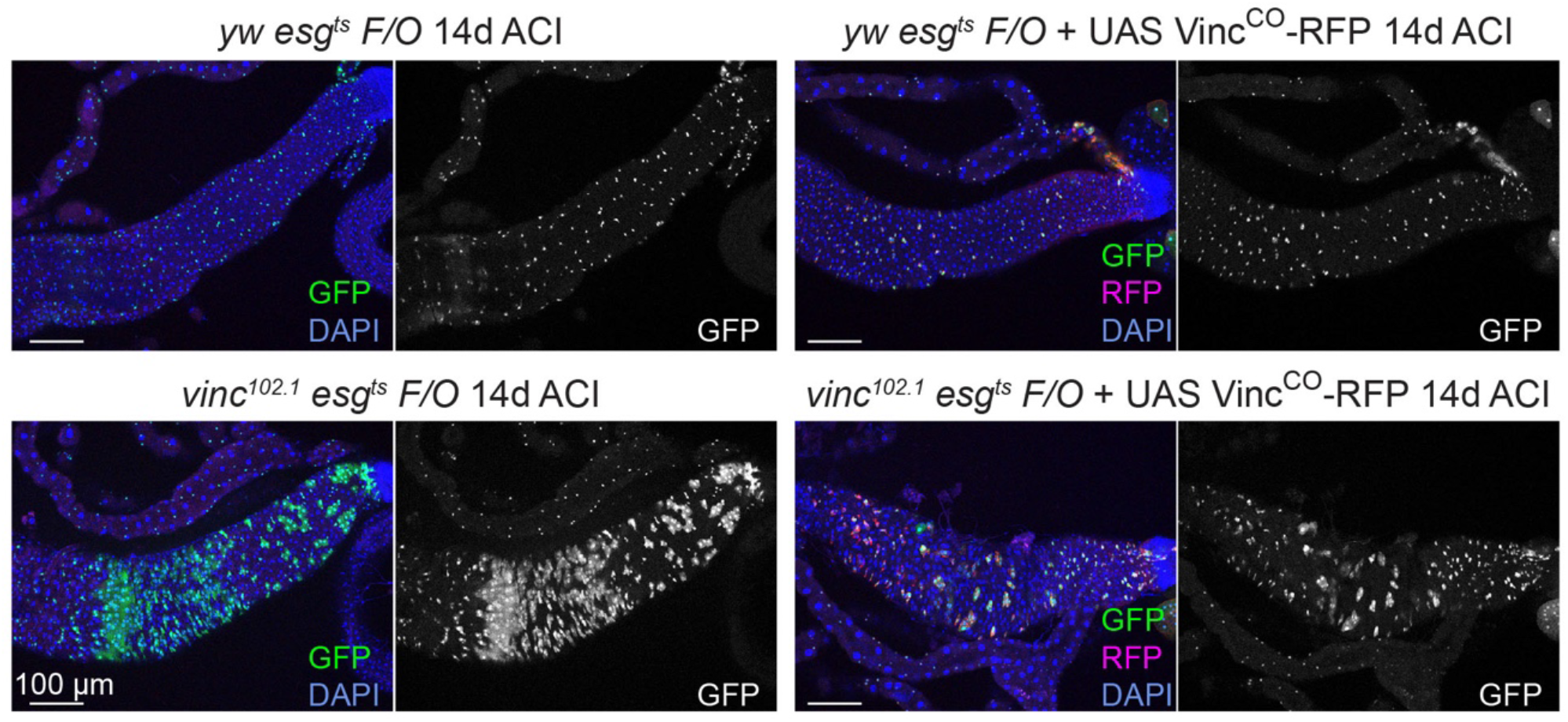
Tissue turnover was assessed in *yw* or *vinc*^*102*.*1*^ guts expressing the *esg*^*ts*^ *F/O* system, with or without *UAS-Vinc*^*CO*^*-RFP* (see fig.2 for methodology). Expression of Vinc^CO^-RFP in progenitors and their newly produced cells (GFP^+^, green) 14 days after clone induction (ACI) prevents the formation of large GFP^+^ patches in the R4/5 region of *vinc*^*102*.*1*^ *esg*^*ts*^ *F/O* guts, indicating that tissue turnover was no longer accelerated as in *vinc*^*102*.*1*^ guts. Expression of Vinc^CO^-RFP in *yw* control guts did not affect turnover rates. GFP (green/white) marks progenitors and newly produced cells, RFP (red) marks cells expressing Vinc^CO^, DAPI (blue) marks nuclei. Images are z-projections over 10μm. Scale bar, 100μm.

**Supplementary Figure 10:**
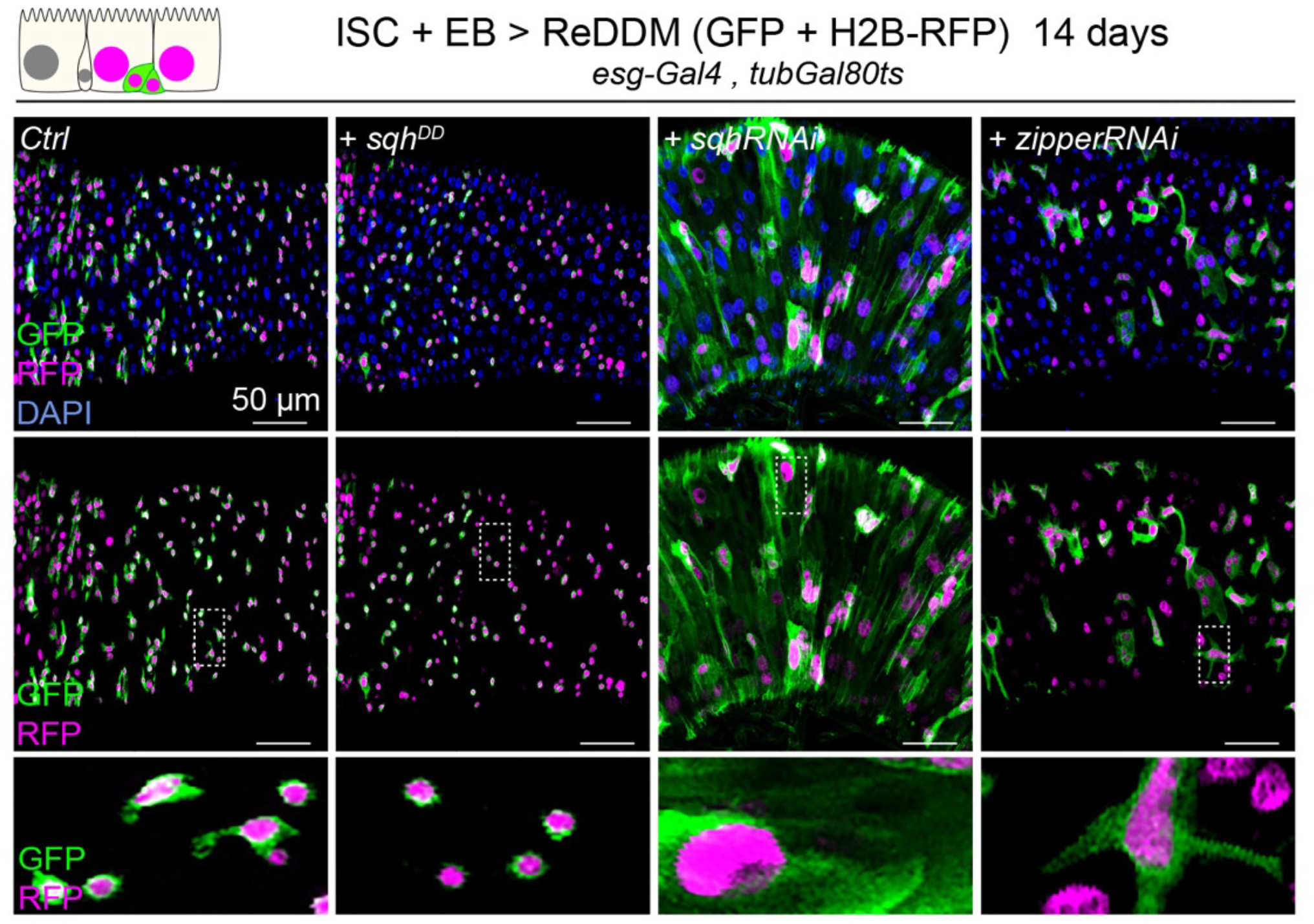
Expression of ReDDM in ISCs and EBs for 14 days with *sqh*^*DD*^, *sqhRNAi* or *zipperRNAi* compared to controls (*Ctrl*). Top panels show the overlay of RFP, GFP and DAPI (nuclei, blue). Middle and bottom panels show RFP and GFP only. Bottom panels show high magnification pictures of the boxes outlined in middle panels. Progenitors round up and remain small upon constitutive activation of Sqh but increase in size and undergo differentiation upon *sqh* or *zipper* knockdown.

**Supplementary Figure 11:**
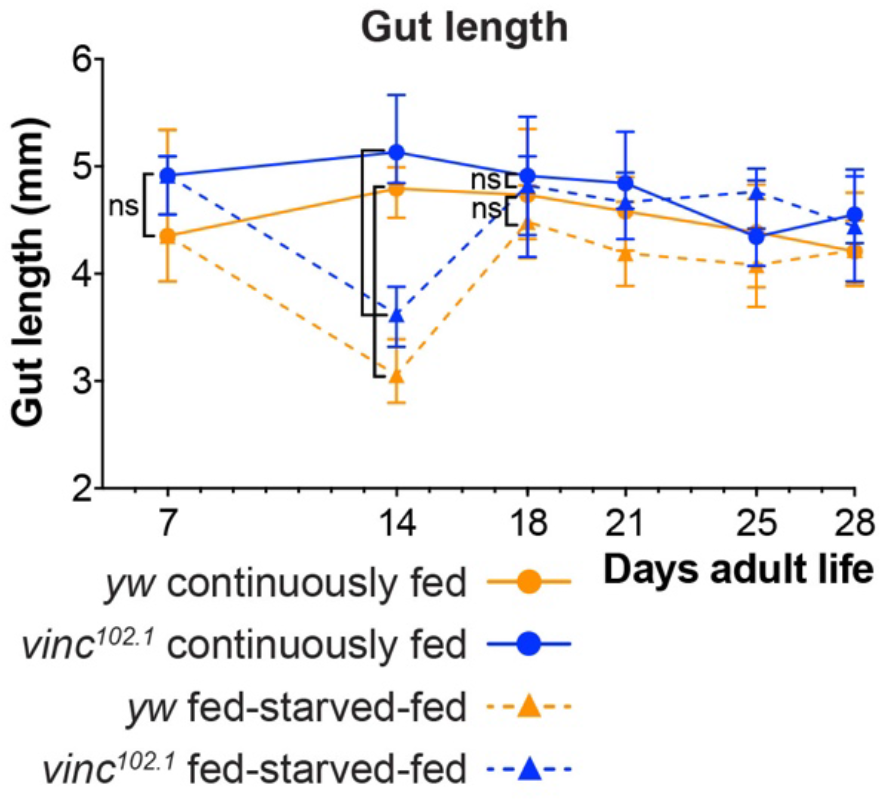
Midgut length changes during cycle of feeding, starving and refeeding. Midgut length is the same in *yw* and *vinc*^*102*.*1*^ after 7 days of feeding (p=0.1782). Following 7 days of starvation, midgut length is significantly shorter in fed-starved compared to continuously fed flies for both *yw* (p<0.0001) and *vinc*^*102*.*1*^ (p<0.0001). Midgut length recovers after just four days of refeeding, with midgut length of fed-starved-refed flies the same as that for continuously fed flies for both *yw* (p=0.1978, ns) and *vinc*^*102*.*1*^ (p=0.6947, ns). This recovery is maintained for the remainder of the experiment. Two-tailed Mann-Whitney test. Symbols represent the median of 9-23 midguts measured, error bars indicate interquartile range.

## References

Alatortsev, V. E., Kramerova, I. A., Frolov, M. V., Lavrov, S. A., & Westphal, E. D. (1997). Vinculin gene is non-essential in Drosophila melanogaster. FEBS Letters, 413(2), 197–201. https://doi.org/10.1016/S0014-5793(97)00901-0

Alégot, H., Markosian, C., Rauskolb, C., Yang, J., Kirichenko, E., Wang, Y. C., & Irvine, K. D. (2019). Recruitment of Jub by α-catenin promotes Yki activity and Drosophila wing growth. Journal of Cell Science, 132(5). https://doi.org/10.1242/jcs.222018

Antonello, Z. a, Reiff, T., Ballesta-Illan, E., & Dominguez, M. (2015). Robust intestinal homeostasis relies on cellular plasticity in enteroblasts mediated by miR-8-Escargot switch. The EMBO Journal, 34(15), 2025– 2041. https://doi.org/10.15252/embj.201591517

Atherton, P., Stutchbury, B., Jethwa, D., & Ballestrem, C. (2016). Mechanosensitive components of integrin adhesions: Role of vinculin. Experimental Cell Research, 343(1), 21–27. https://doi.org/10.1016/j.yexcr.2015.11.017

Bach, E. A., Ekas, L. A., Ayala-Camargo, A., Flaherty, M. S., Lee, H., Perrimon, N., & Baeg, G. H. (2007). GFP reporters detect the activation of the Drosophila JAK/STAT pathway in vivo. Gene Expression Patterns, 7(3), 323–331. https://doi.org/10.1016/j.modgep.2006.08.003

Bays, J. L., & DeMali, K. A. (2017). Vinculin in cell–cell and cell–matrix adhesions. Cellular and Molecular Life Sciences, 74(16), 2999–3009. https://doi.org/10.1007/s00018-017-2511-3

Beebe, K., Lee, W. C., & Micchelli, C. A. (2010). JAK/STAT signaling coordinates stem cell proliferation and multilineage differentiation in the Drosophila intestinal stem cell lineage. Developmental Biology, 338(1), 28–37. https://doi.org/10.1016/j.ydbio.2009.10.045

Bharadwaj, R., Roy, M., Ohyama, T., Sivan-Loukianova, E., Delannoy, M., Lloyd, T. E., Zlatic, M., Eberl, D. F., & Kolodkin, A. L. (2013). Cbl-associated protein regulates assembly and function of two tension-sensing structures in Drosophila. Development (Cambridge), 140(3), 627–638. https://doi.org/10.1242/dev.085100

Biswas, R., Banerjee, A., Lembo, S., Zhao, Z., Lakshmanan, V., Lim, R., Le, S., Nakasaki, M., Kutyavin, V., Wright, G., Palakodeti, D., Ross, R. S., Jamora, C., Vasioukhin, V., Jie, Y., & Raghavan, S. (2021). Mechanical instability of adherens junctions overrides intrinsic quiescence of hair follicle stem cells. Developmental Cell, 56(6), 761-780.e7. https://doi.org/10.1016/j.devcel.2021.02.020

Buchon, N., Broderick, N. A., Kuraishi, T., & Lemaitre, B. (2010). Drosophila EGFR pathway coordinates stem cell proliferation and gut remodeling following infection. BMC Biology, 8. https://doi.org/10.1186/1741-7007-8-152

Buchon, N., Osman, D., David, F. P. A., Yu Fang, H., Boquete, J. P., Deplancke, B., & Lemaitre, B. (2013). Morphological and molecular characterization of adult midgut compartmentalization in Drosophila. Cell Reports, 3(5), 1725–1738. https://doi.org/10.1016/j.celrep.2013.04.001

Carisey, A., & Ballestrem, C. (2011). Vinculin, an adapter protein in control of cell adhesion signalling. European Journal of Cell Biology, 90(2–3), 157–163. https://doi.org/10.1016/j.ejcb.2010.06.007

Case, L. B., Baird, M. A., Shtengel, G., Campbell, S. L., Hess, H. F., Davidson, M. W., & Waterman, C. M. (2015). Molecular mechanism of vinculin activation and nanoscale spatial organization in focal adhesions. Nature Cell Biology, 17(7), 880–892. https://doi.org/10.1038/ncb3180

Chen, J., Sayadian, A. C., Lowe, N., Lovegrove, H. E., & St Johnston, D. (2018). An alternative mode of epithelial polarity in the Drosophila midgut. PLoS Biology, 16(10), 1–24. https://doi.org/10.1371/journal.pbio.3000041

Choi, N. H., Lucchetta, E., & Ohlstein, B. (2011). Nonautonomous regulation of Drosophila midgut stem cell proliferation by the insulin-signaling pathway. Proceedings of the National Academy of Sciences of the United States of America, 108(46), 18702–18707. https://doi.org/10.1073/pnas.1109348108

De Navascués, J., Perdigoto, C. N., Bian, Y., Schneider, M. H., Bardin, A. J., Martínez-Arias, A., & Simons, B. D. (2012). Drosophila midgut homeostasis involves neutral competition between symmetrically dividing intestinal stem cells. EMBO Journal, 31(11), 2473–2485. https://doi.org/10.1038/emboj.2012.106

Del Rio, A., Perez-Jimenez, R., Liu, R., Roca-Cusachs, P., Fernandez, J. M., & Sheetz, M. P. (2009). Stretching single talin rod molecules activates vinculin binding. Science, 323(5914), 638–641. https://doi.org/10.1126/science.1162912

Dumbauld, D. W., Lee, T. T., Singh, A., Scrimgeour, J., Gersbach, C. A., Zamir, E. A., Fu, J., Chen, C. S., Curtis, J. E., Craig, S. W., & García, A. J. (2013). How vinculin regulates force transmission. Proceedings of the National Academy of Sciences of the United States of America, 110(24), 9788–9793. https://doi.org/10.1073/pnas.1216209110

Franke, J. D., Boury, A. L., Gerald, N. J., & Kiehart, D. P. (2006). Native nonmuscle myosin II stability and light chain binding in Drosophila melanogaster. Cell Motility and the Cytoskeleton, 63(10), 604–622. https://doi.org/10.1002/cm.20148

Furriols, M., & Bray, S. (2001). A model Notch response element detects suppressor of hairless-dependent molecular switch. Current Biology, 11(1), 60–64. https://doi.org/10.1016/S0960-9822(00)00044-0

Gehart, H., & Clevers, H. (2019). Tales from the crypt: new insights into intestinal stem cells. Nature Reviews Gastroenterology and Hepatology, 16(1), 19–34. https://doi.org/10.1038/s41575-018-0081-y

Gjorevski, N., Sachs, N., Manfrin, A., Giger, S., Bragina, M. E., Ordóñez-Morán, P., Clevers, H., & Lutolf, M. P. (2016). Designer matrices for intestinal stem cell and organoid culture. Nature, 539(7630), 560–564. https://doi.org/10.1038/nature20168

Goldmann, W. H., Auernheimer, V., Thievessen, I., & Fabry, B. (2013). Vinculin, cell mechanics and tumour cell invasion. Cell Biology International, 37(5), 397–405. https://doi.org/10.1002/cbin.10064

Goulas, S., Conder, R., & Knoblich, J. A. (2012). The Par complex and Integrins direct asymmetric cell division in adult intestinal stem cells. Cell Stem Cell, 11(4), 529–540. https://doi.org/10.1016/j.stem.2012.06.017

Green, H. J., Griffiths, A. G. M., Ylänne, J., & Brown, N. H. (2018). Novel functions for integrin-associated proteins revealed by analysis of myofibril attachment in Drosophila. ELife, 7, 1–29. https://doi.org/10.7554/eLife.35783

Guisoni, N., Martinez-Corral, R., Garcia-Ojalvo, J., & de Navascués, J. (2017). Diversity of fate outcomes in cell pairs under lateral inhibition. Development (Cambridge), 144(7), 1177–1186. https://doi.org/10.1242/dev.137950

He, L., Si, G., Huang, J., Samuel, A. D. T., & Perrimon, N. (2018). Mechanical regulation of stem-cell differentiation by the stretch-activated Piezo channel. Nature. https://doi.org/10.1038/nature25744

Hermiston, M. L., & Gordon, J. I. (1995). In vivo analysis of cadherin function in the mouse intestinal epithelium: Essential roles in adhesion, maintenance of differentiation, and regulation of programmed cell death. Journal of Cell Biology, 129(2), 489–506. https://doi.org/10.1083/jcb.129.2.489

Hermiston, M. L., Wong, M. H., Gordon, J. I., Wong, M. H., Hermiston, M. L., Syder, A. J., & Gordon, J. I. (1999). Forced expression of E-cadherin in the mouse intestinal epithelium slows cell migration and provides evidence for nonautonomous regulation of cell fate in a self-renewing system: Forced expression of the tumor suppressor adenomatosis polyposis coli protei. Chemtracts, 12(3), 200–209.

Holle, A. W., Tang, X., Vijayraghavan, D., Vincent, L. G., Fuhrmann, A., Choi, Y. S., Del Alamo, J. C., & Engler, Á.J. (2013). In situ mechanotransduction via vinculin regulates stem cell differentiation. Stem Cells, 31(11), 2467–2477. https://doi.org/10.1002/stem.1490

Hu, D. J. K., & Jasper, H. (2019). Control of intestinal cell fate by dynamic mitotic spindle repositioning influences epithelial homeostasis and longevity. Cell Reports, 28(11), 2807-2823.e5. https://doi.org/10.1016/j.celrep.2019.08.014

Hynes, R. (2002). Integrins: bidirectional, allosteric signaling machines. Cell, 110(6), 673–687. https://doi.org/10.1016/s0092-8674(02)00971-6.

Jiang, H., Patel, P. H., Kohlmaier, A., Grenley, M. O., McEwen, D. G., & Edgar, B. A. (2009). Cytokine/Jak/Stat Signaling mediates regeneration and homeostasis in the Drosophila midgut. Cell, 137(7), 1343–1355. https://doi.org/10.1016/j.cell.2009.05.014

Jurado, J., Navascués, J.de, & Gorfinkiel, N. (2016). α-Catenin stabilises Cadherin-Catenin complexes and modulates actomyosin dynamics to allow pulsatile apical contraction. Journal of Cell Science, 129(24), 4496–4508. https://doi.org/10.1242/jcs.193268

Kale, G. R., Yang, X., Philippe, J. M., Mani, M., Lenne, P. F., & Lecuit, T. (2018). Distinct contributions of tensile and shear stress on E-cadherin levels during morphogenesis. Nature Communications, 9(1). https://doi.org/10.1038/s41467-018-07448-8

Klapholz, B., & Brown, N. H. (2017). Talin - The master of integrin adhesions. Journal of Cell Science, 130(15), 2435–2446. https://doi.org/10.1242/jcs.190991

Klapholz, B., Herbert, S. L., Wellmann, J., Johnson, R., Parsons, M., & Brown, N. H. (2015). Alternative mechanisms for talin to mediate integrin function. Current Biology, 25(7), 847–857. https://doi.org/10.1016/j.cub.2015.01.043

Kohlmaier, A., Fassnacht, C., Jin, Y., Reuter, H., Begum, J., Dutta, D., & Edgar, B. A. (2015). Src kinase function controls progenitor cell pools during regeneration and tumor onset in the Drosophila intestine. Oncogene, 34(18), 2371–2384. https://doi.org/10.1038/onc.2014.163

Korzelius, J., Azami, S., Ronnen-Oron, T., Koch, P., Baldauf, M., Meier, E., Rodriguez-Fernandez, I. A., Groth, M., Sousa-Victor, P., & Jasper, H. (2019). The WT1-like transcription factor Klumpfuss maintains lineage commitment of enterocyte progenitors in the Drosophila intestine. Nature Communications, 10(1). https://doi.org/10.1038/s41467-019-12003-0

Krndija, D., Marjou FEl, Guirao, B., Richon, S., Leroy, O., Bellaiche, Y., Hannezo, E., & Vignjevic, D. M. (2019). Active cell migration is critical for steady-state epithelial turnover in the gut. Science, 365(6454), 705–710. https://doi.org/10.1126/science.aau3429

Kumar, A., Placone, J. K., & Engler, A. J. (2017). Understanding the extracellular forces that determine cell fate and maintenance. Development (Cambridge), 144(23), 4261–4270. https://doi.org/10.1242/dev.158469

Kuroda, M., Wada, H., Kimura, Y., Ueda, K., & Kioka, N. (2017). Vinculin promotes nuclear localization of TAZ to inhibit ECM stiffness-dependent differentiation into adipocytes. Journal of Cell Science, 130(5), 989–1002. https://doi.org/10.1242/jcs.194779

Le Duc, Q., Shi, Q., Blonk, I., Sonnenberg, A., Wang, N., Leckband, D., & De Rooij, J. (2010). Vinculin potentiates E-cadherin mechanosensing and is recruited to actin-anchored sites within adherens junctions in a myosin II-dependent manner. Journal of Cell Biology, 189(7), 1107–1115. https://doi.org/10.1083/jcb.201001149

Li, Q., Nirala, N. K., Nie, Y., Chen, H. J., Ostroff, G., Mao, J., Wang, Q., Xu, L., & Ip, Y. T. (2018). Ingestion of food particles regulates the mechanosensing Misshapen-Yorkie pathway in Drosophila intestinal growth. Developmental Cell, 45(4), 433-449.e6. https://doi.org/10.1016/j.devcel.2018.04.014

Li, T., Guo, H., Song, Y., Zhao, X., Shi, Y., Lu, Y., Hu, S., Nie, Y., Fan, D., & Wu, K. (2014). Loss of vinculin and membrane-bound β-catenin promotes metastasis and predicts poor prognosis in colorectal cancer. Molecular Cancer, 13(1), 1–15. https://doi.org/10.1186/1476-4598-13-263

Liang, J., Balachandra, S., Ngo, S., & O’Brien, L. E. (2017). Feedback regulation of steady-state epithelial turnover and organ size. Nature, 548(7669), 588–591. https://doi.org/10.1038/nature23678

Lin, G., Zhang, X., Ren, J., Pang, Z., Wang, C., Xu, N., & Xi, R. (2013a). Integrin signaling is required for maintenance and proliferation of intestinal stem cells in Drosophila. Developmental Biology, 377(1), 177– 187. https://doi.org/10.1016/j.ydbio.2013.01.032

Lin, G., Zhang, X., Ren, J., Pang, Z., Wang, C., Xu, N., & Xi, R. (2013b). Integrin signaling is required for maintenance and proliferation of intestinal stem cells in Drosophila. Developmental Biology, 377(1), 177– 187. https://doi.org/10.1016/j.ydbio.2013.01.032

Lucchetta, E. M., & Ohlstein, B. (2017). Amitosis of polyploid cells regenerates functional stem cells in the Drosophila intestine. Cell Stem Cell, 20(5), 609-620.e6. https://doi.org/10.1016/j.stem.2017.02.012

Maartens, A. P., Wellmann, J., Wictome, E., Klapholz, B., Green, H., & Brown, N. H. (2016). Drosophila vinculin is more harmful when hyperactive than absent, and can circumvent integrin to form adhesion complexes. Journal of Cell Science, 129(23), 4354–4365. https://doi.org/10.1242/jcs.189878

Maeda, K., Takemura, M., Umemori, M., & Adachi-Yamada, T. (2008). E-cadherin prolongs the moment for interaction between intestinal stem cell and its progenitor cell to ensure Notch signaling in adult Drosophila midgut. Genes to Cells, 13(12), 1219–1227. https://doi.org/10.1111/j.1365-2443.2008.01239.x

Marianes, A., & Spradling, A. C. (2013). Physiological and stem cell compartmentalization within the Drosophila midgut. ELife, 2013(2), 1–19. https://doi.org/10.7554/eLife.00886

Martin, J. L., Sanders, E. N., Moreno-Roman, P., Koyama, L. A. J., Balachandra, S., Du, X., & O’brien, L. E. (2018). Long-term live imaging of the Drosophila adult midgut reveals real-time dynamics of division, differentiation and loss. ELife, 7, 1–33. https://doi.org/10.7554/eLife.36248

McLeod, C. J., Wang, L., Wong, C., & Jones, D. L. (2010). Stem cell dynamics in response to nutrient availability. Current Biology, 20(23), 2100–2105. https://doi.org/10.1016/j.cub.2010.10.038

Meran, L., Baulies, A., & Li, V. S. W. (2017). Intestinal stem cell niche: the extracellular matrix and cellular components. Stem Cells International, 2017. https://doi.org/10.1155/2017/7970385

Mesa, K. R., Kawaguchi, K., Cockburn, K., Gonzalez, D., Boucher, J., Xin, T., Klein, A. M., & Greco, V. (2018). Homeostatic epidermal stem cell self-renewal is driven by local differentiation. Cell Stem Cell, 23(5), 677-686.e4. https://doi.org/10.1016/j.stem.2018.09.005

Micchelli, C. A., & Perrimon, N. (2006). Evidence that stem cells reside in the adult Drosophila midgut epithelium. Nature, 439(7075), 475–479. https://doi.org/10.1038/nature04371

Miguel-Aliaga, I., Jasper, H., & Lemaitre, B. (2018). Anatomy and physiology of the digestive tract of Drosophila melanogaster. Genetics, 210(2), 357–396. https://doi.org/10.1534/genetics.118.300224

Mitonaka, T., Muramatsu, Y., Sugiyama, S., Mizuno, T., & Nishida, Y. (2007). Essential roles of myosin phosphatase in the maintenance of epithelial cell integrity of Drosophila imaginal disc cells. Developmental Biology, 309(1), 78–86. https://doi.org/10.1016/j.ydbio.2007.06.021

Monster JL, Donker L, Vliem MJ, Win Z, Matthews HK, Cheah JS, Yamada S, de Rooij J, Baum B, Gloerich M. (2021) An asymmetric junctional mechanoresponse coordinates mitotic rounding with epithelial integrity. J Cell Biol. 220(5):e202001042. doi: 10.1083/jcb.202001042

O’Brien, L. E., Soliman, S. S., Li, X., & Bilder, D. (2011). Altered modes of stem cell division drive adaptive intestinal growth. Cell, 147(3), 603–614. https://doi.org/10.1016/j.cell.2011.08.048

Ohlstein, B., & Spradling, A. (2006). The adult Drosophila posterior midgut is maintained by pluripotent stem cells. Nature, 439(7075), 470–474. https://doi.org/10.1038/nature04333

Ohlstein, B., & Spradling, A. (2007). Multipotent Drosophila intestinal stem cells specify daughter cell fates by differential notch signaling. Science, 315(5814), 988–992. https://doi.org/10.1126/science.1136606

Okumura, T., Takeda, K., Taniguchi, K., & Adachi-Yamada, T. (2014). β? Integrin inhibits chronic and high level activation of JNK to repress senescence phenotypes in Drosophila adult midgut. PLoS ONE, 9(2), e89387. https://doi.org/10.1371/journal.pone.0089387

Reiff, T., Antonello, Z. A., Ballesta-Illán, E., Mira, L., Sala, S., Navarro, M., Martinez, L. M., & Dominguez, M. (2019). Notch and EGFR regulate apoptosis in progenitor cells to ensure gut homeostasis in Drosophila . The EMBO Journal, 38(21), 1–15. https://doi.org/10.15252/embj.2018101346

Rezaie, A., Park, S. C., Morales, W., Marsh, E., Lembo, A., Kim, J. H., Weitsman, S., Chua, K. S., Barlow, G. M., & Pimentel, M. (2017). Assessment of anti-vinculin and anti-cytolethal distending Toxin B antibodies in subtypes of irritable bowel syndrome. Digestive Diseases and Sciences, 62(6), 1480–1485. https://doi.org/10.1007/s10620-017-4585-z

Rojas Villa, S. E., Meng, F. W., & Biteau, B. (2019). Zfh2 controls progenitor cell activation and differentiation in the adult Drosophila intestinal absorptive lineage. PLoS Genetics, 15(12), 1–24. https://doi.org/10.1371/journal.pgen.1008553

Rübsam, M., Mertz, A. F., Kubo, A., Marg, S., Jüngst, C., Goranci-Buzhala, G., Schauss, A. C., Horsley, V., Dufresne, E. R., Moser, M., Ziegler, W., Amagai, M., Wickström, S. A., & Niessen, C. M. (2017). E-cadherin integrates mechanotransduction and EGFR signaling to control junctional tissue polarization and tight junction positioning. Nature Communications, 8(1), 1–15. https://doi.org/10.1038/s41467-017-01170-7

Sarpal, R., Yan, V., Kazakova, L., Sheppard, L., & Tepass, U. (2019). Role of α-Catenin and its mechanosensing properties in the regulation of Hippo/YAP-dependent tissue growth. PLoS Genetics, 1–30. https://doi.org/https://doi.org/10.1371/journal.pgen.1008454

Sun, Z., Guo, S. S., & Fässler, R. (2016). Integrin-mediated mechanotransduction. J Cell Biol. 215(4):445–456. doi: 10.1083/jcb.201609037.

Thomas, W. A., Boscher, C., Chu, Y. S., Cuvelier, D., Martinez-Rico, C., Seddiki, R., Heysch, J., Ladoux, B., Thiery, J. P., Mege, R. M., & Dufour, S. (2013). α-Catenin and vinculin cooperate to promote high E-cadherin-based adhesion strength. Journal of Biological Chemistry, 288(7), 4957–4969. https://doi.org/10.1074/jbc.M112.403774

Wang, L., Zeng, X., Ryoo, H. D., & Jasper, H. (2014). Integration of UPRER and oxidative stress signaling in the control of intestinal stem cell proliferation. PLoS Genetics, 10(8). https://doi.org/10.1371/journal.pgen.1004568

Wang, P., Wu, J., Wood, A., Jones, M., Pedley, R., Li, W., Ross, R. S., Ballestrem, C., Gilmore, A. P., & Streuli, C. H. (2019). Vinculins interaction with talin is essential for mammary epithelial differentiation. Scientific Reports, 9(1), 18400. https://doi.org/10.1038/s41598-019-54784-w

Wickström, S. A., & Niessen, C. M. (2018). Cell adhesion and mechanics as drivers of tissue organization and differentiation: local cues for large scale organization. Current Opinion in Cell Biology, 54, 89–97. https://doi.org/10.1016/j.ceb.2018.05.003

Xiang, J., Bandura, J., Zhang, P., Jin, Y., Reuter, H., & Edgar, B. A. (2017). EGFR-dependent TOR-independent endocycles support Drosophila gut epithelial regeneration. Nature Communications, 8(May), 1–13. https://doi.org/10.1038/ncomms15125

Xu, W., Baribault, H., & Adamson, E. D. (1998). Vinculin knockout results in heart and brain defects during embryonic development. Development, 125(2), 327–337.

Yao, M., Qiu, W., Liu, R., Efremov, A. K., Cong, P., Seddiki, R., Payre, M., Lim, C. T., Ladoux, B., Mège, R. M., & Yan, J. (2014). Force-dependent conformational switch of α-catenin controls vinculin binding. Nature Communications, 5. https://doi.org/10.1038/ncomms5525

Yonemura, S., Wada, Y., Watanabe, T., Nagafuchi, A., & Shibata, M. (2010). α-Catenin as a tension transducer that induces adherens junction development. Nature Cell Biology, 12(6), 533–542. https://doi.org/10.1038/ncb2055

You, J., Zhang, Y., Li, Z., Lou, Z., Jin, L., & Lin, X. (2014). Drosophila Perlecan regulates intestinal stem cell activity via cell-matrix attachment. Stem Cell Reports, 2(6), 761–769. https://doi.org/10.1016/j.stemcr.2014.04.007

Zhai, Z., Boquete, J. P., & Lemaitre, B. (2017). A genetic framework controlling the differentiation of intestinal stem cells during regeneration in Drosophila. PLoS Genetics, 13(6), 1–27. https://doi.org/10.1371/journal.pgen.1006854

